## Supplementary Material for "A new deep-branching environmental lineage of algae"

### New deep-branching environmental plastid genomes on the algal tree of life

**This pdf file contains**

**Supplementary Figure 1-26**
**Supplementary Tables 1-12**
**Supplementary Note 1**

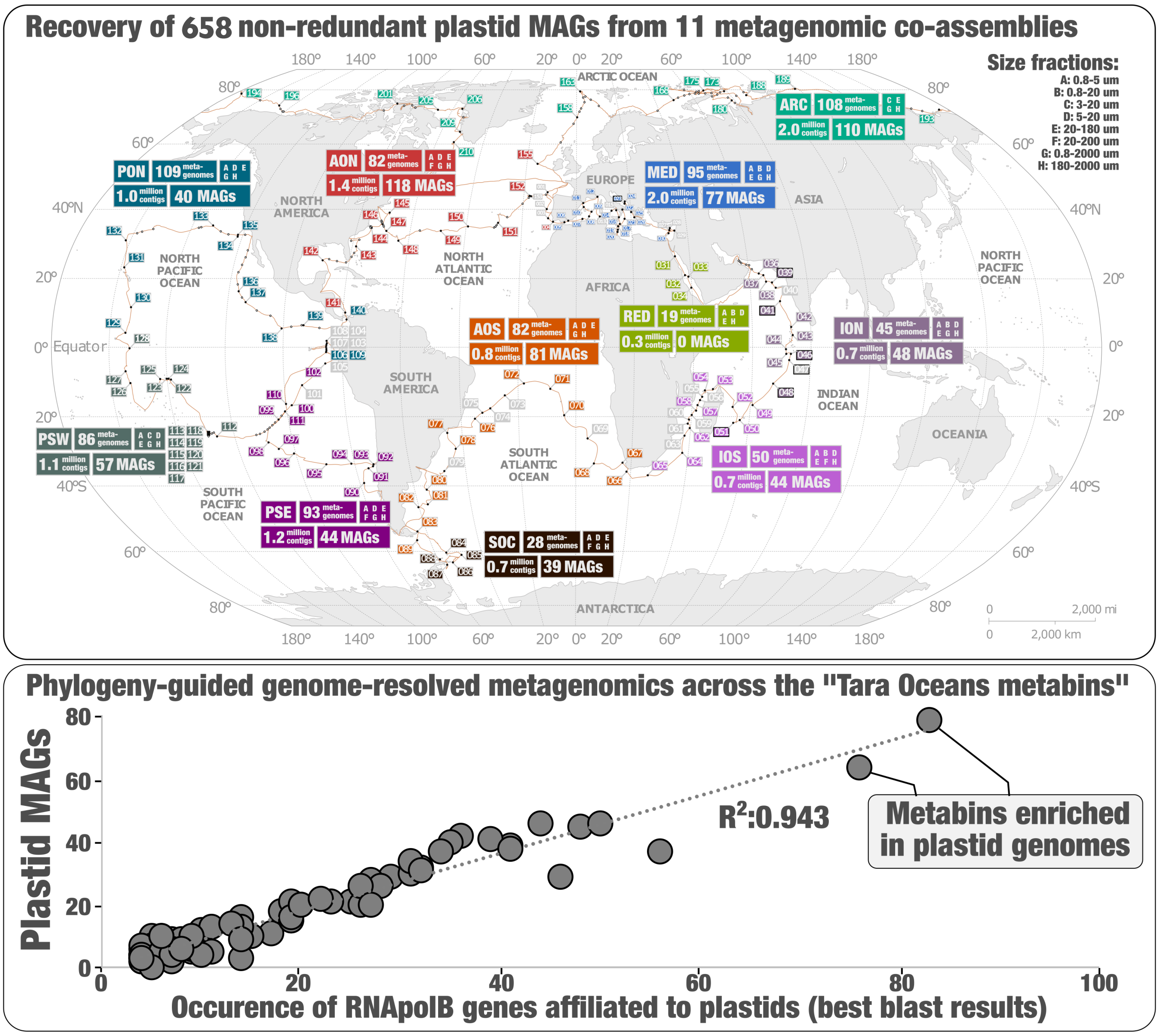


**Supplementary Fig. 1.** Top panel summarizes how many non-redundant ptMAGs were characterized from each of the 11 large *Tara* Oceans co-assemblies. Bottom panel shows the strong correlation between (1) the number of plastid-affiliated RNApolB genes occurring in each *Tara* Oceans metabin (each metabin contains a subset of contigs from one co-assembly that were automatically binned using differential coverage and sequence composition), (2) and the number of manually characterized ptMAGs across the same metabins.

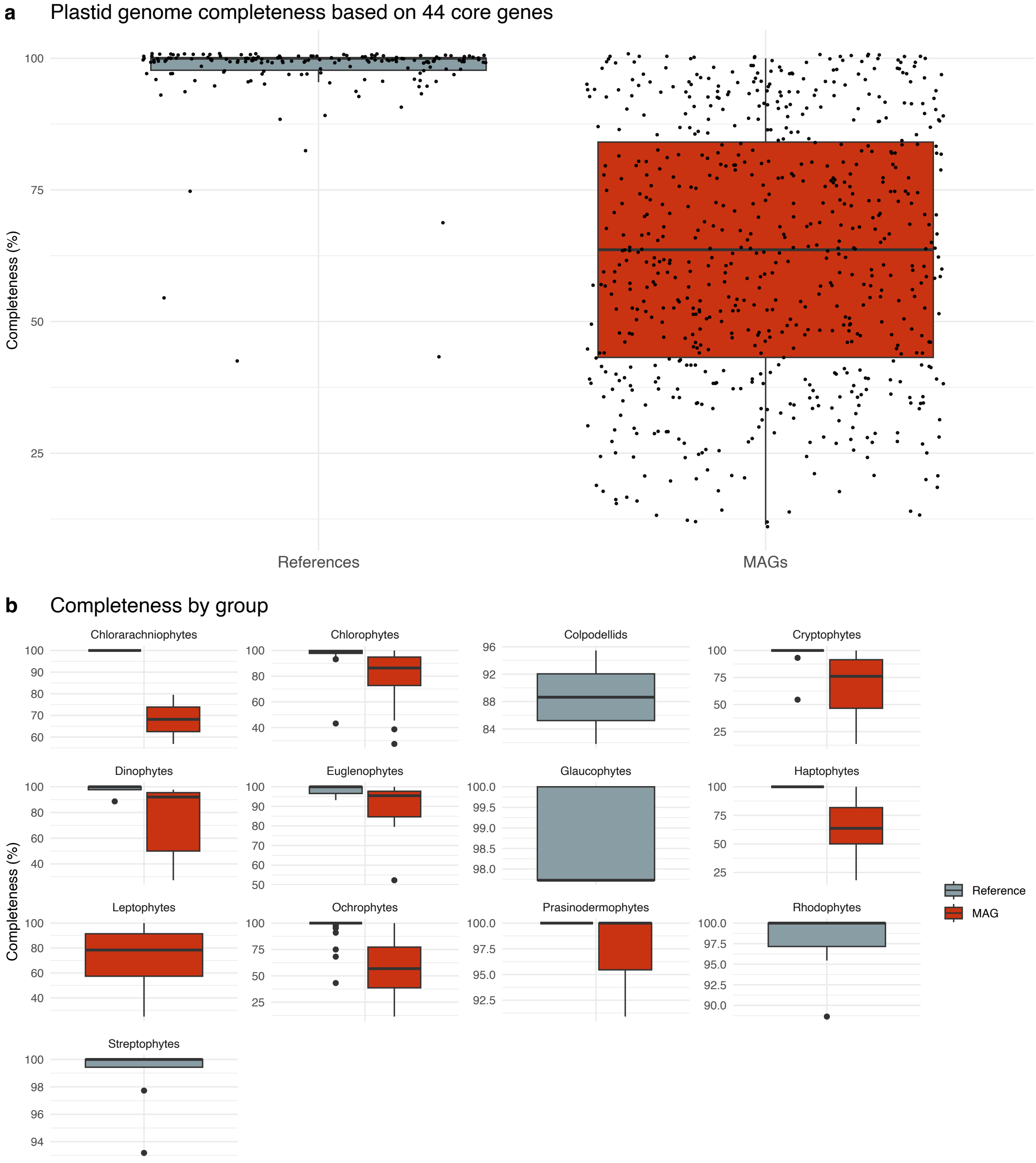


**Supplementary Fig. 2.** Estimated completeness of the reference plastid genomes (n=165, not including cyanobacteria and *Paulinella*) and ptMAGs (n=660). Panel (a) displays an overall comparison between references and ptMAGs with dots representing individual genomes, while panel (b) presents a comparison between groups with dots representing outlier genomes. Completeness was estimated based on the presence of 44 core genes (listed in Supplementary Table 4).

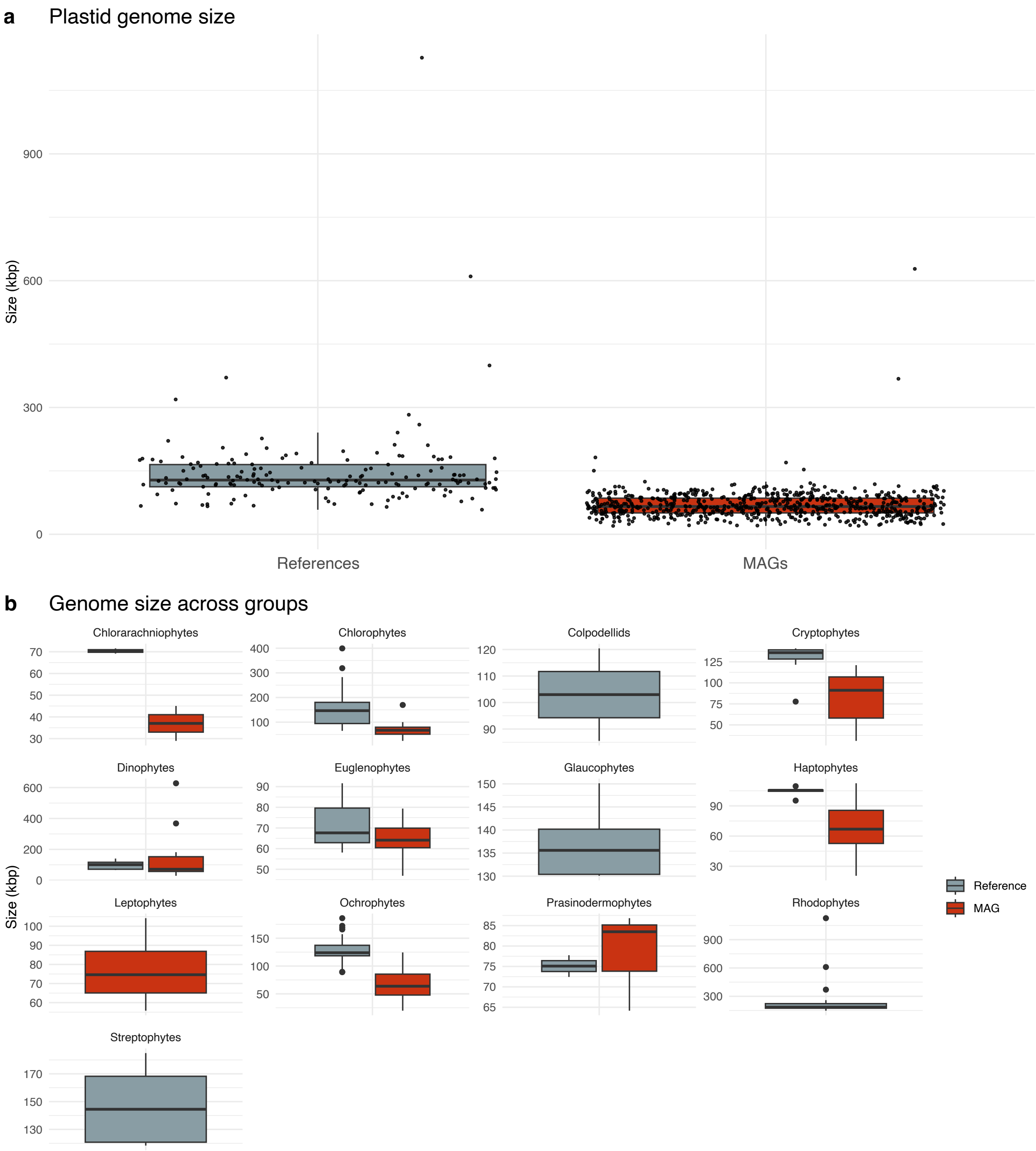


**Supplementary Fig. 3.** Size of the reference plastid genomes (n=165, not including cyanobacteria and *Paulinella*) and ptMAGs (n=660). Panel (a) shows that ptMAGs are smaller than reference genomes in general, with dots representing individual genomes, while panel (b) presents a comparison between groups with dots representing outlier genomes.

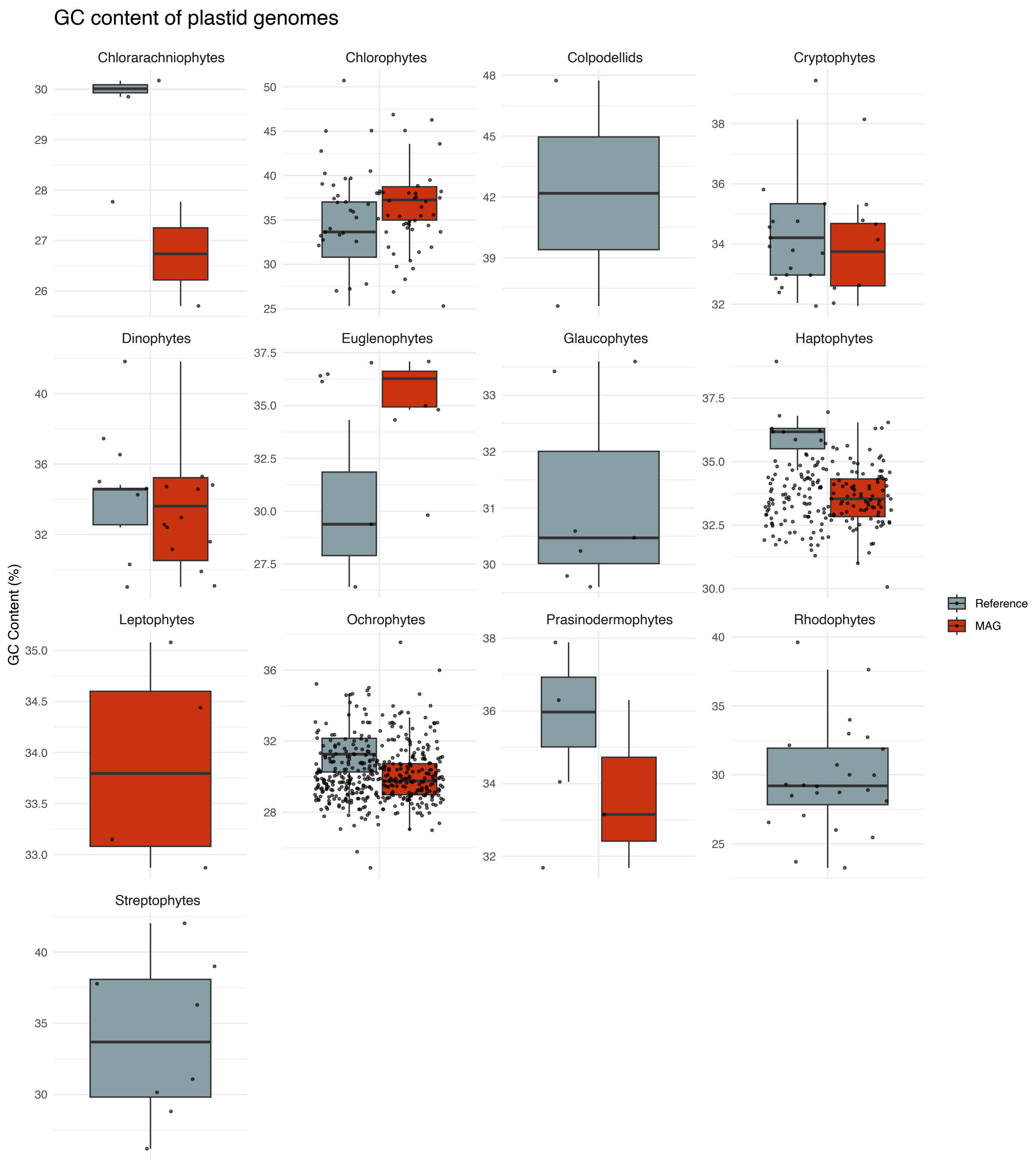


**Supplementary Fig. 4.** GC content of the plastid genomes across algal groups including references (n=165, not including cyanobacteria and *Paulinella*) and ptMAGs (n=660).

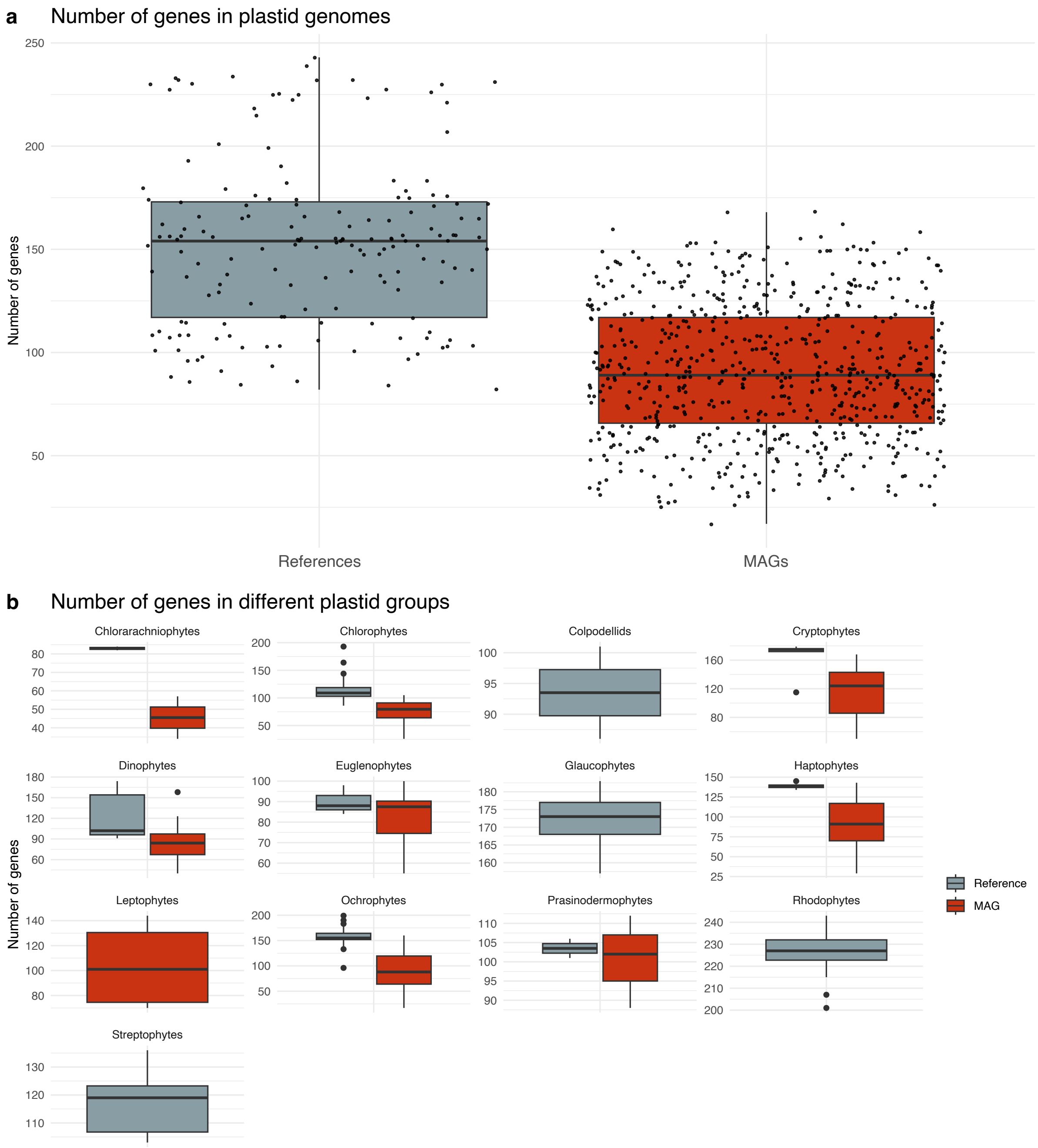


**Supplementary Fig. 5.** Number of genes encoded in the reference plastid genomes (n=165, not including cyanobacteria and *Paulinella*) and ptMAGs (n=660) included in the Maximum Likelihood phylogeny shown in Figure 1. Panel (a) shows that ptMAGs generally encode fewer genes due to lower completeness than reference genomes, with dots representing individual genomes, while panel (b) presents a comparison between groups with dots representing outlier genomes.

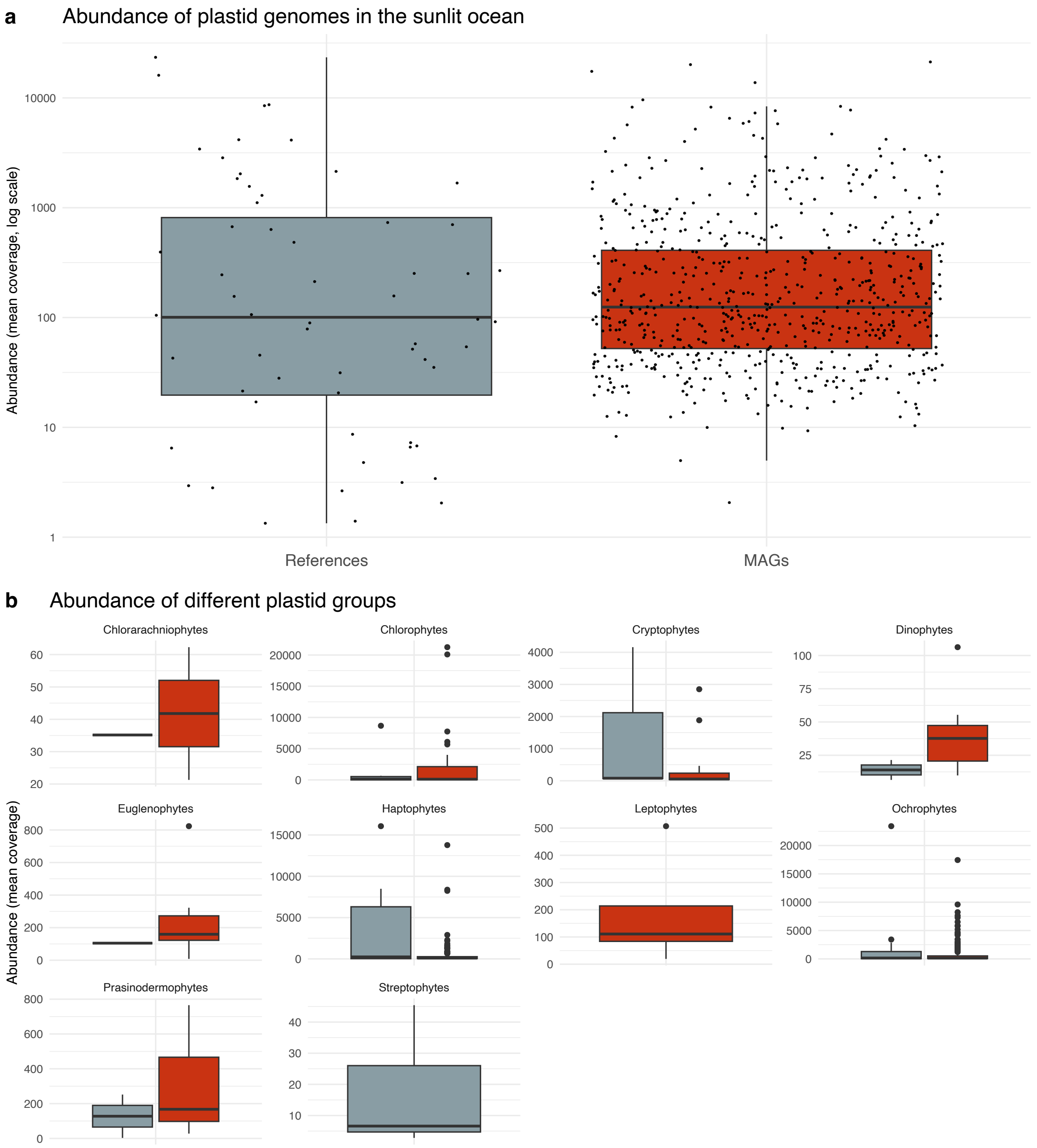


**Supplementary Fig. 6.** Abundance of plastid genomes in the sunlit ocean as estimated by mean vertical coverage obtained by mapping 937 *Tara* Oceans metagenomes to each genome. To account for non-specific read recruitment, a genome was only considered to be detected if at least 25% of its length was covered by reads. Panel (a) and (b) are based on 60 reference genomes (not including cyanobacteria, *Paulinella*, and non-marine taxa) and 660 ptMAGs. ptMAGs represent nearly 85% of the total signal, in part due to the low number of selected reference genomes, and in many cases, references still represent the most abundant taxa (e.g., ochrophytes, haptophytes, and cryptophytes).

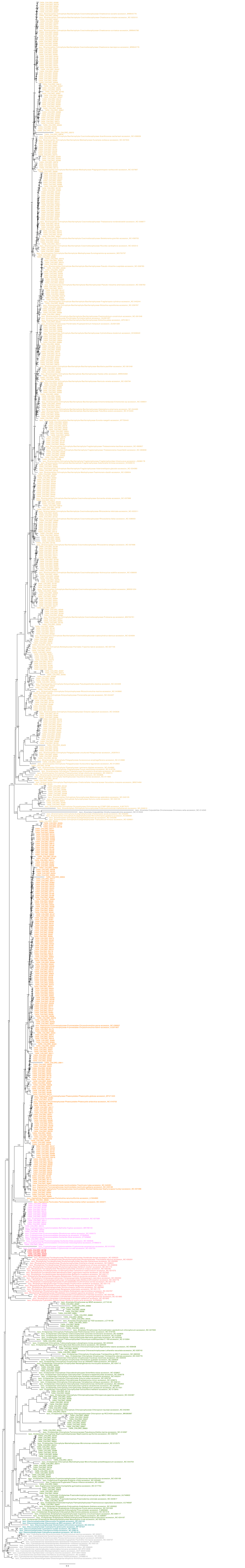

**Supplementary Fig. 7.** (On previous page). Maximum-likelihood phylogeny of 179 selected reference genomes and 660 ptMAGs based on 93 plastid-encoded genes. The tree was run under the LG+F+I+G4 model, and branch support values were calculated with 1000 ultrafast bootstrap replicates.

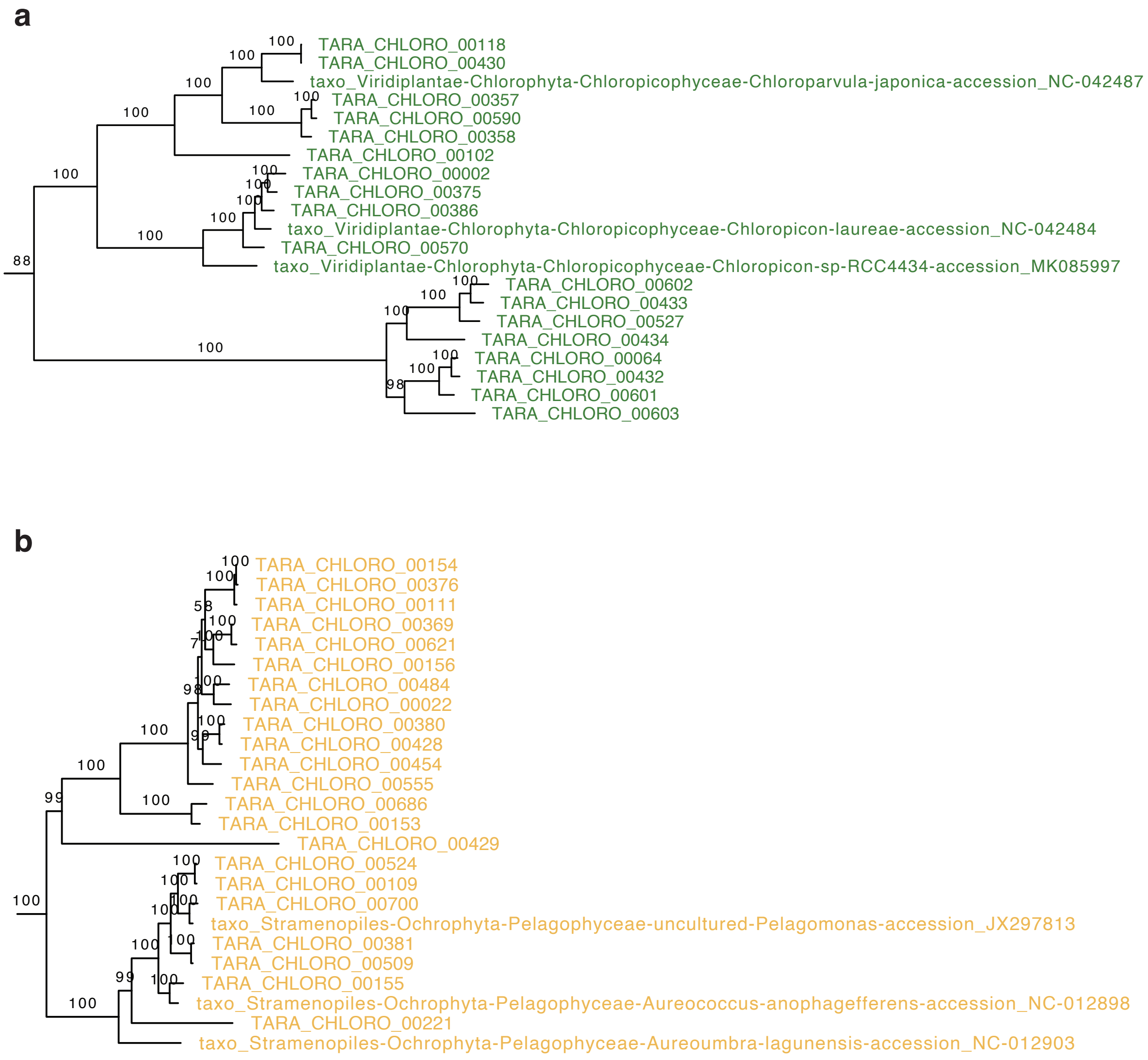


**Supplementary Fig. 8.** Two examples of deep-branching novel diversity in algal groups detected by ptMAGs. Panel (a) shows a clade of eight ptMAGs that is related to Chloropicophyceae. Panel (b) shows a novel clade of plastid genomes that is related to, but distinct from known pelagophyte references.

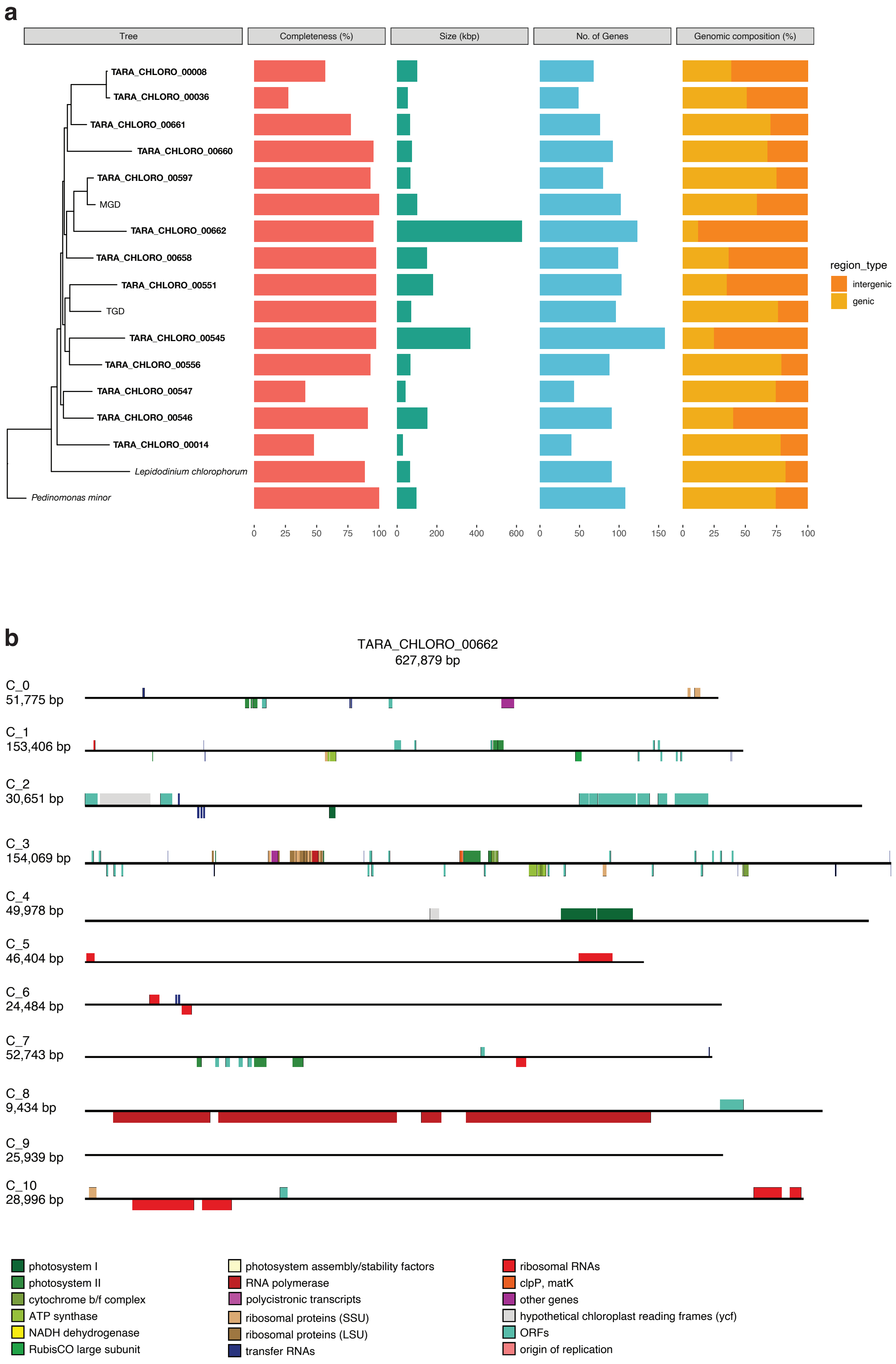

a
Tree
Completeness (%)
Size (kbp)
No. of Genes
Genomic composition (%)
TARA_CHLORO_00008
TARA_CHLORO_00036
TARA_CHLORO_00661
TARA_CHLORO_00660
TARA_CHLORO_00597
MGD
TARA_CHLORO_00662
TARA_CHLORO_00658
TARA_CHLORO_00551
TGD
TARA_CHLORO_00545
TARA_CHLORO_00556
TARA_CHLORO_00547
TARA_CHLORO_00546
TARA_CHLORO_00014
Lepidodinium chlorophorum
Pedinomonas minor
0 25 50 75 100
0 200 400 600
0 50 100 150
0 25 50 75 100
region_type
intergenic
genic
b
TARA_CHLORO_00662
627,879 bp
C_0
51,775 bp
C_1
153,406 bp
C_2
30,651 bp
C_3
154,069 bp
C_4
49,978 bp
C_5
46,404 bp
C_6
24,484 bp
C_7
52,743 bp
C_8
9,434 bp
C_9
25,939 bp
C_10
28,996 bp
photosystem I
photosystem II
cytochrome b/f complex
ATP synthase
NADH dehydrogenase
RubisCO large subunit
photosystem assembly/stability factors
RNA polymerase
polycistronic transcripts
ribosomal proteins (SSU)
ribosomal proteins (LSU)
transfer RNAs
ribosomal RNAs
clpP, matK
other genes
hypothetical chloroplast reading frames (ycf)
ORFs
origin of replication

**Supplementary Fig. 9.** (on previous page) **(a)** A clade of 13 ptMAGs related to green algal endosymbionts of dinoflagellates, showing large variations in size, with the largest genome (TARA_CHLORO_00662) being around 600 kbp. This variation in genome size is accompanied by an inflation in intergenic regions in the plastid genome. Panel **(b)** visualises the plastome organisation (including large intergenic regions) in the largest ptMAG. Contigs are not drawn to scale and size is indicated.

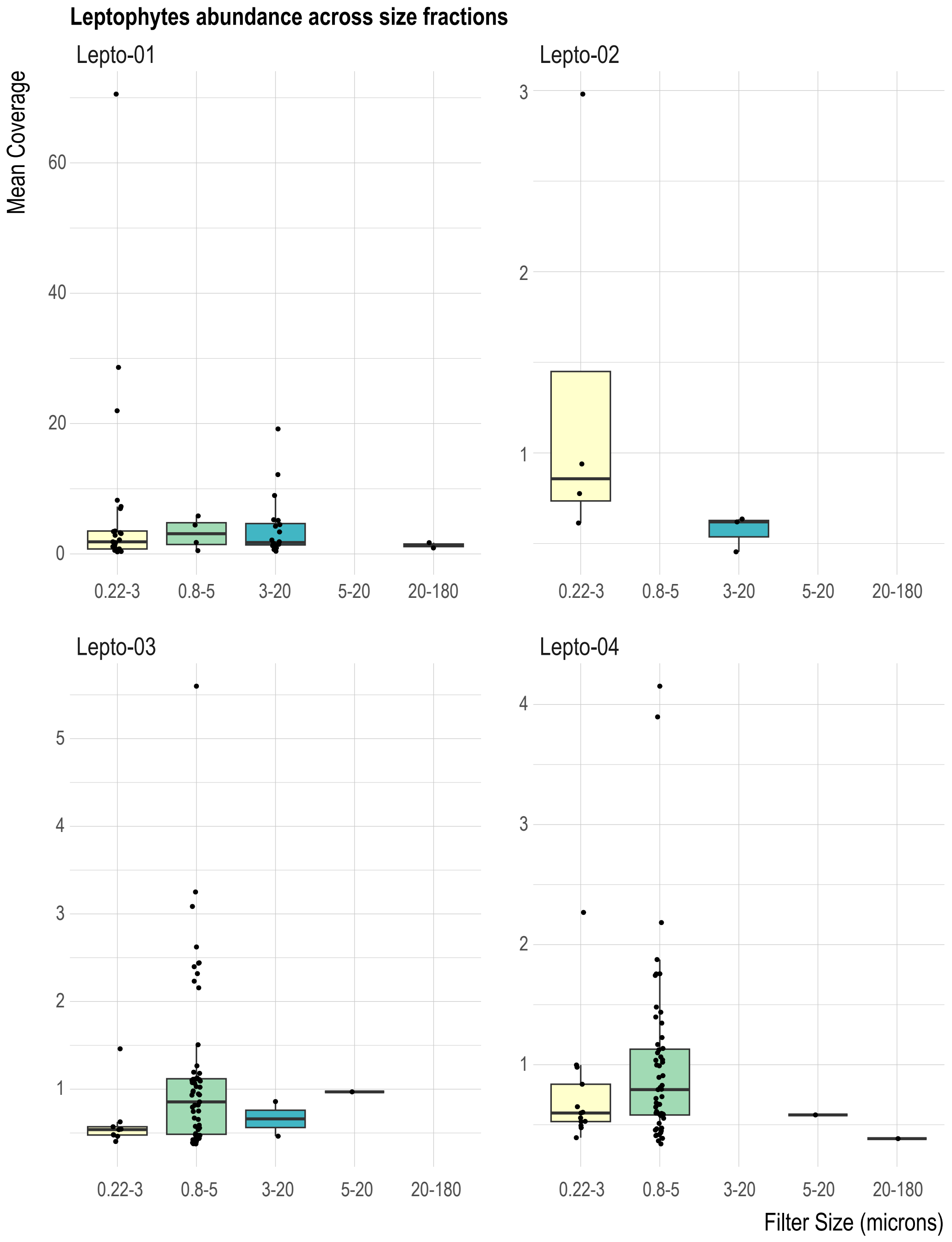


**Supplementary Fig. 10.** Signal of leptophyte plastid genomes in *Tara* Ocean samples from different size fractions. Leptophytes are detected most frequently in the 0.22-3 $\mu$m, 0.8-5 $\mu$m, and the 3-20 $\mu$m size fractions, and are barely detected in the 5-20 $\mu$m and 20-180 $\mu$m size fractions, indicating that leptophytes are small algae that are less than 5 $\mu$m in size.

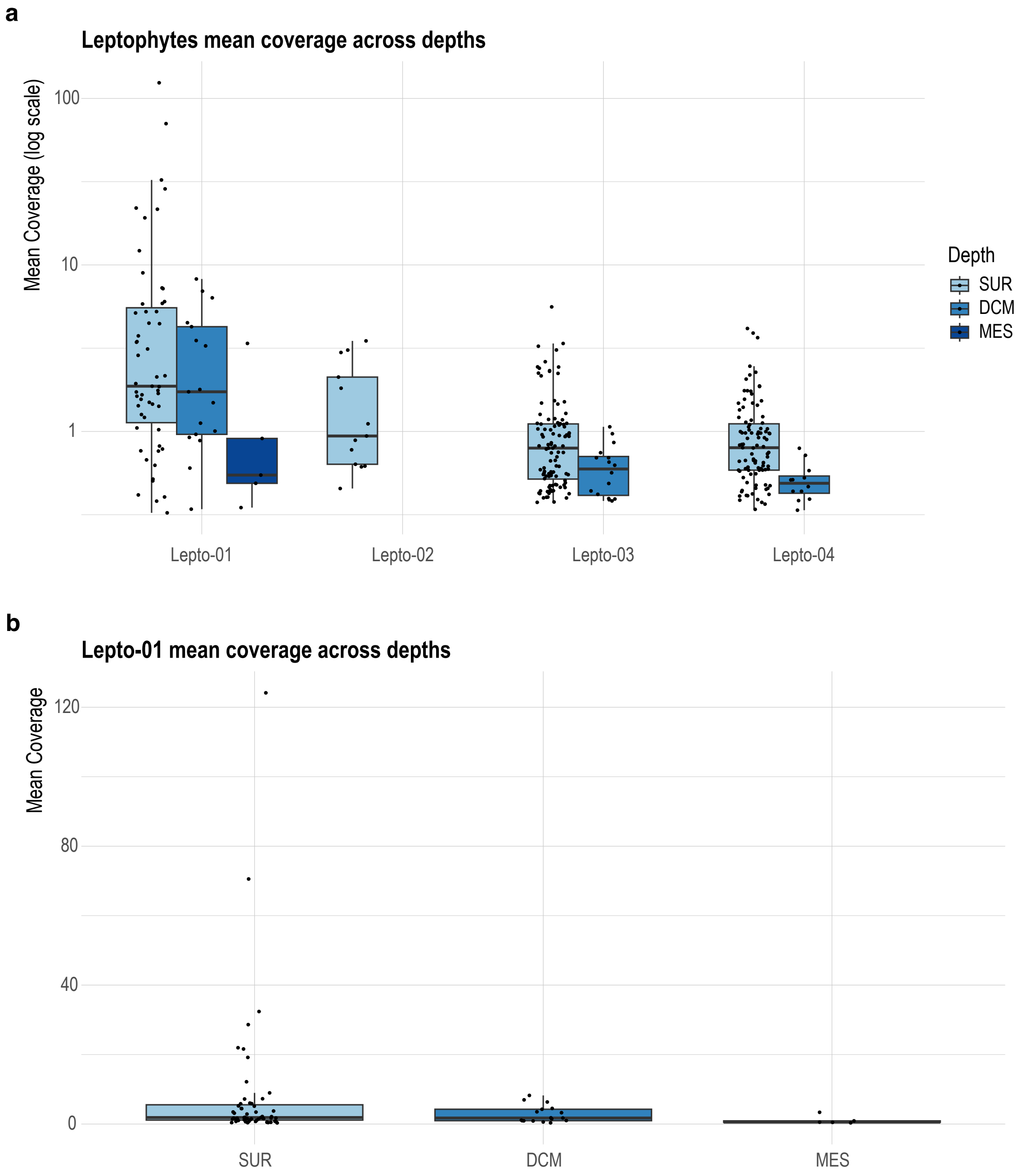


**Supplementary Fig. 11.** Signal of leptophyte plastid genomes in *Tara* Ocean samples from different depths. Panel **(a)** depicts all four leptophyte ptMAGs while panel **(b)** zooms in on the most abundant ptMAG (Lepto-01). Data points represent mean coverage values from individual samples. Leptophytes are predominantly detected in the surface layer, detected at a lesser abundance in the DCM layer, and are barely detected in the mesopelagic layer.

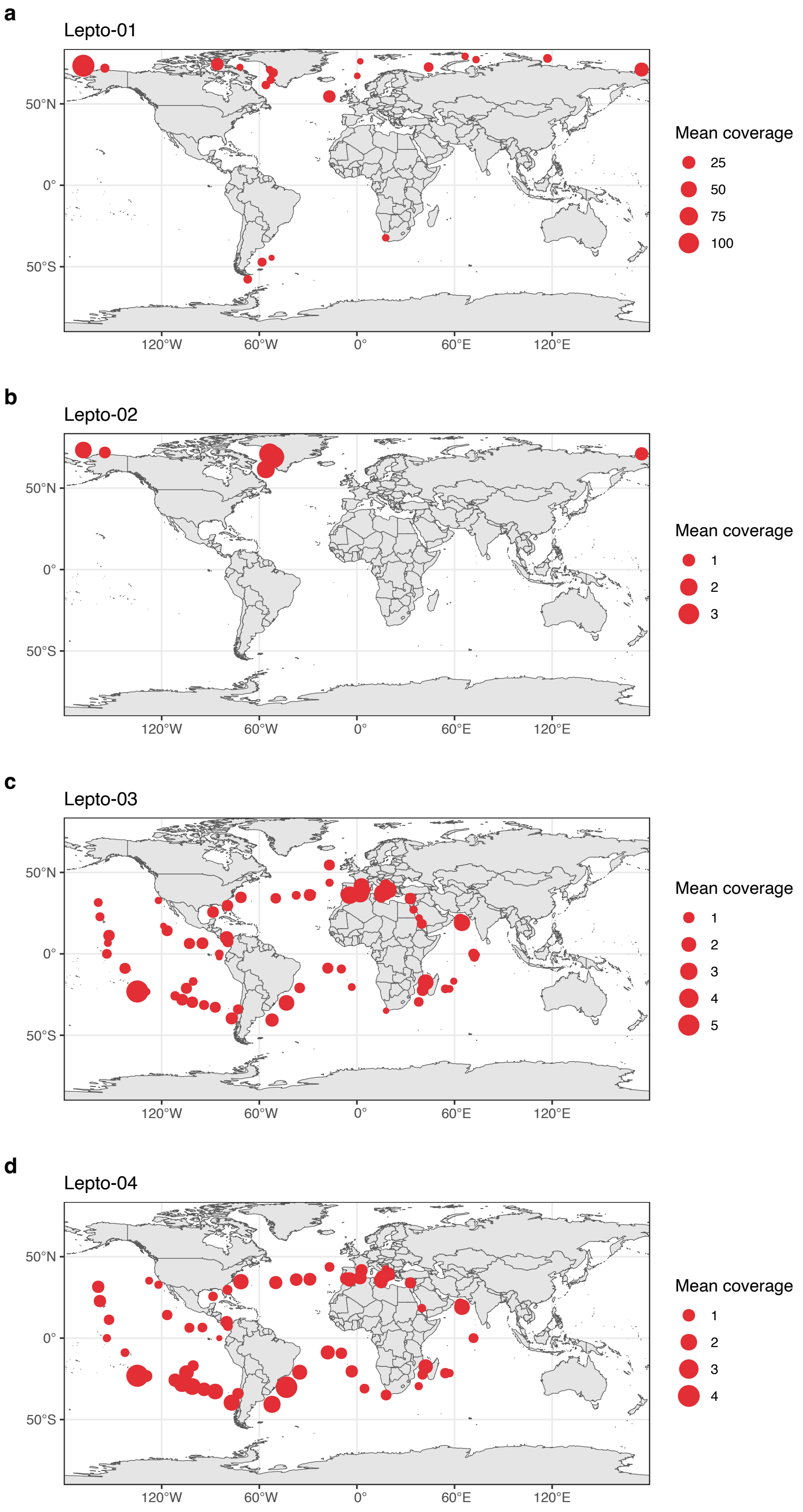


**Supplementary Fig. 12.** Distribution of leptophyte plastid genomes as determined by mapping of 937 *Tara* Ocean metagenomes against each genome. Lepto-01 and Lepto-02 display an Arctic distribution, while Lepto-03 and Lepto-04 ptMAGs have a more tropical and subtropical distribution. The overall low mean coverage indicates that leptophytes are rare in general, with the exception of several stations where Lepto-01 is abundant.

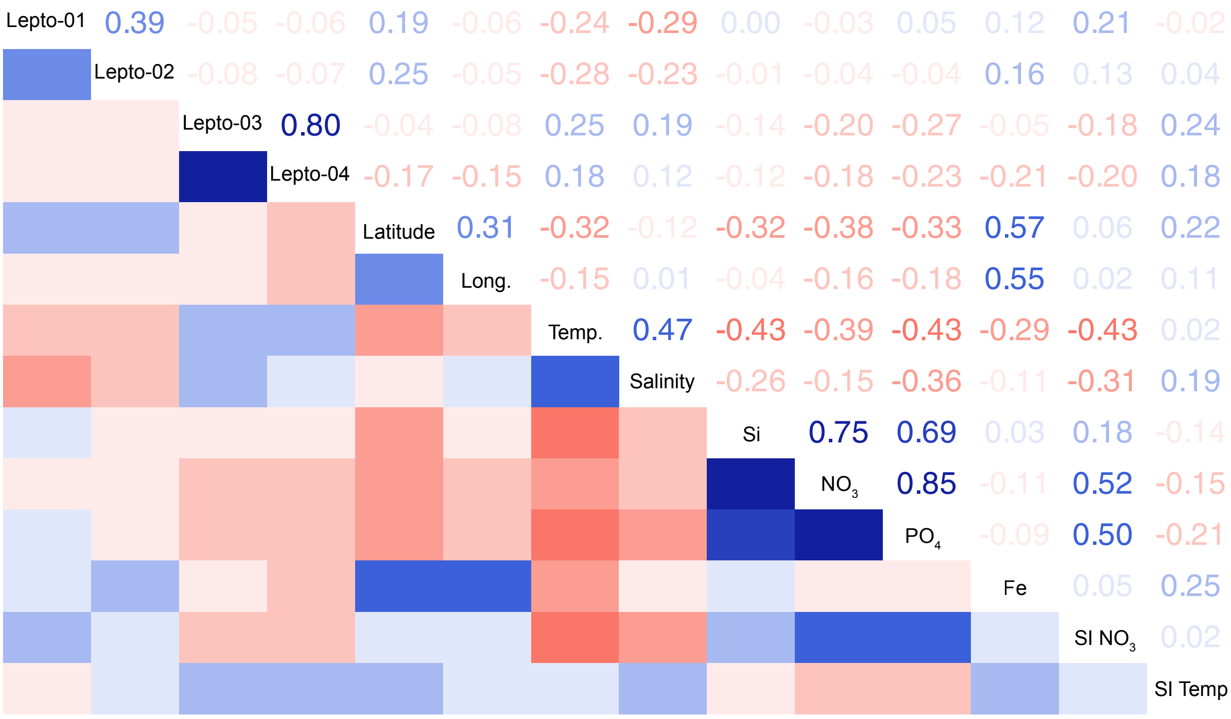


**Supplementary Fig. 13.** Pearson correlations between the abundance of leptophyte plastids and environmental parameters. SI $NO_3$ and SI Temp represent the seasonality indices of nitrate and sea surface temperature respectively. These were defined as the range of the nitrate and temperature in one grid cell divided by the total range of that variable across all *Tara* sampling stations.

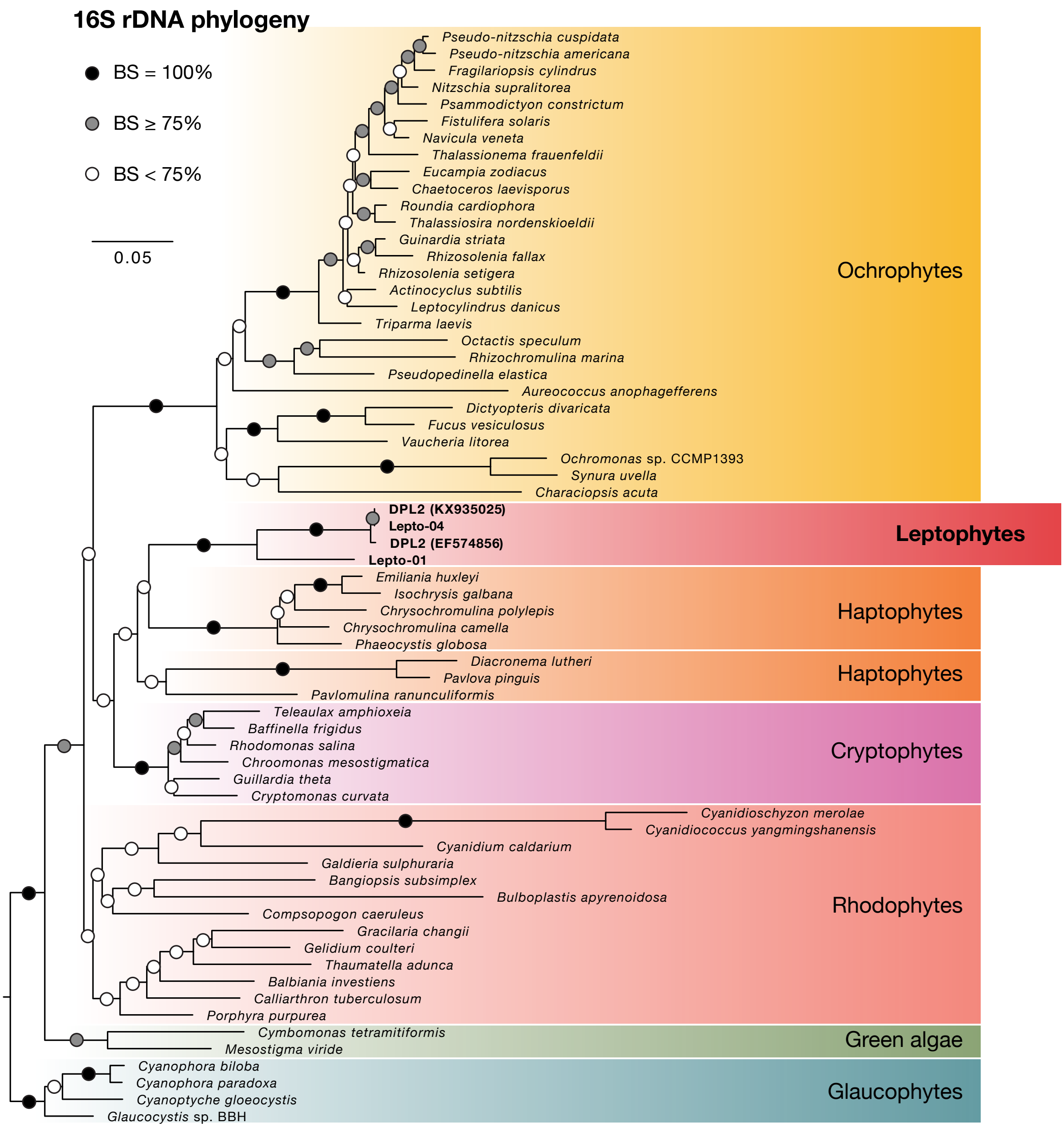


**Supplementary Fig. 14.** Maximum Likelihood phylogeny inferred with an alignment of the plastid 16S rDNA gene (65 taxa, 1499 sites). The phylogeny was inferred with raxml-ng using the GTR+G model and branch support assessed with 100 non-parametric bootstrap replicates.

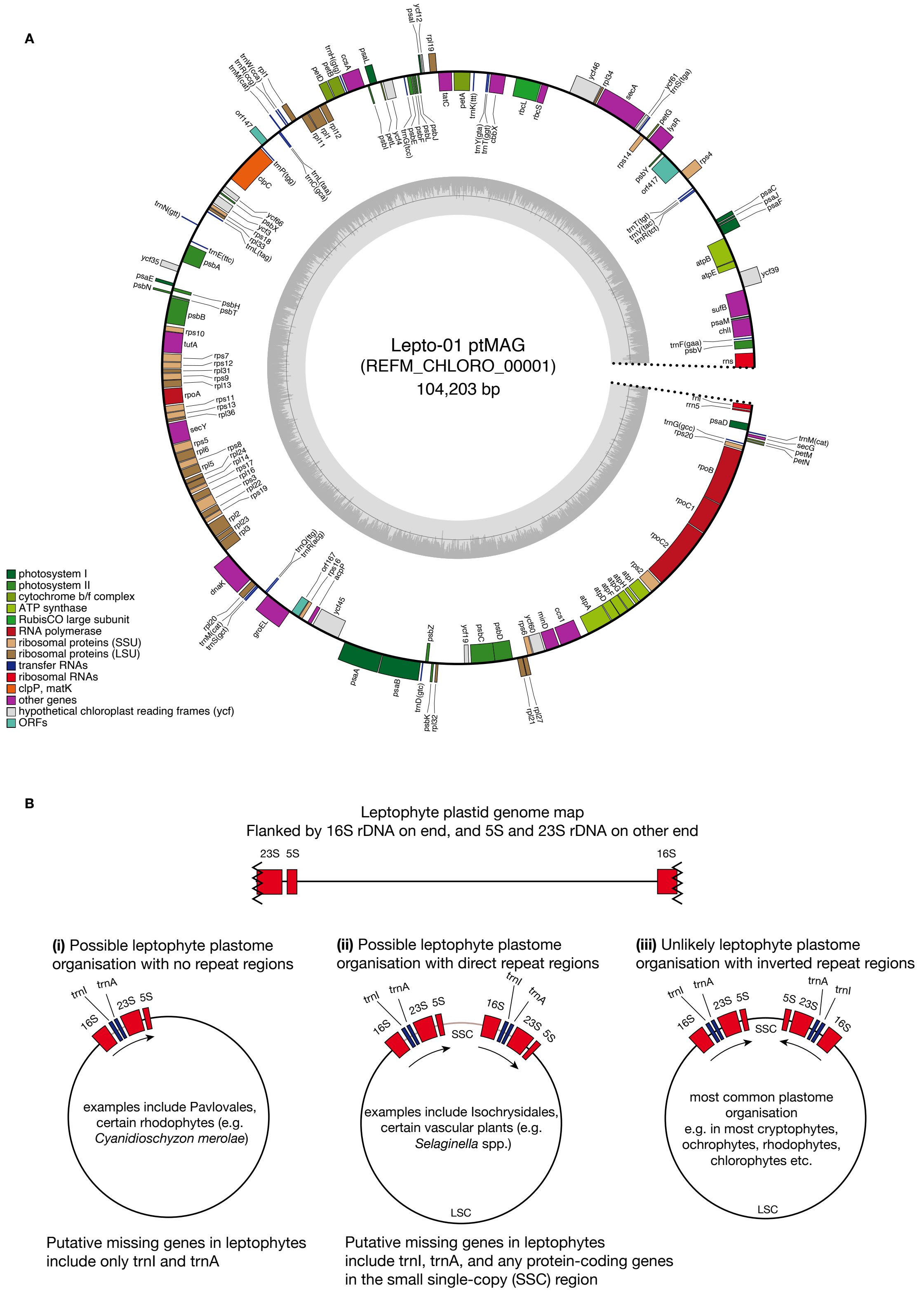


**Supplementary Fig. 15.** **(A)** Annotated map of the Lepto-01 plastid genome. Genes drawn inside the outer circle are transcribed clockwise, while those drawn outside the outer circle are transcribed counterclockwise. Genes are colour coded based on functional groups. The dark shaded area in the inner circle indicates GC content. The genome map was generated by OGDRAW[1]. **(B)** Cartoons of possible plastome organisation of leptophytes based on the available leptophyte plastid genome map.

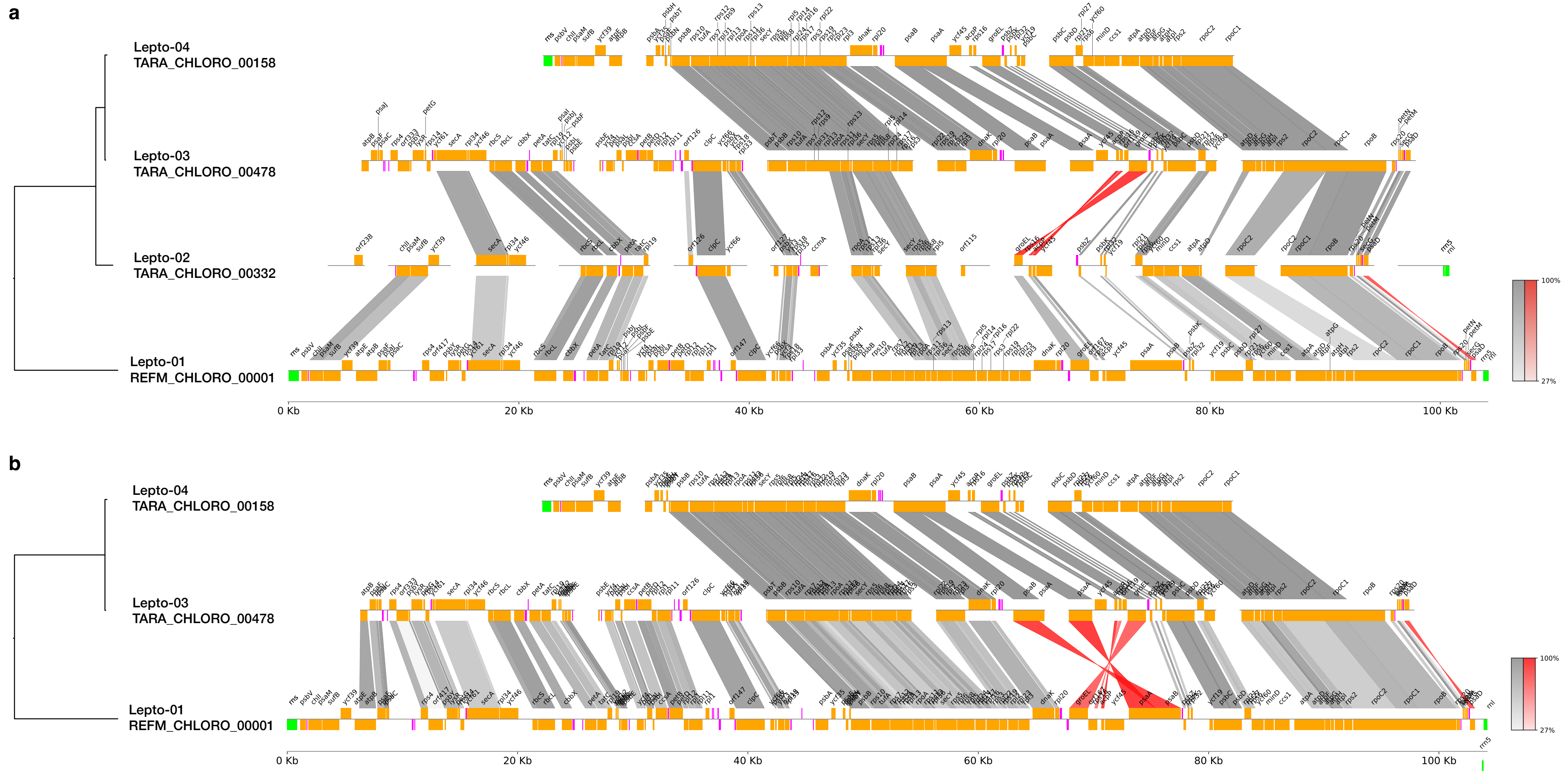


**Supplementary Fig. 16.** Synteny comparisons of the leptophyte ptMAGs. Panel (**a**) displays all leptophyte ptMAGs, and panel (**b**) displays the three most complete ptMAGs. Genomes were visualized using PyGenomeViz and sequence similarity was calculated using MMSEQs for reciprocal best-hit CDS search[2]. A cartoon phylogeny on the left depicts the relationships between these plastid genomes. Protein coding genes are coloured orange and labelled, while tRNA sequences are coloured magenta and not labelled for clarity. Overall, the leptophyte plastid genomes are very similar in gene content and synteny with the exception of two differences: (1) The *psaD* gene is inverted in Lepto-01 compared to the rest; and (2) A nearly 10 kbp region spanning from the *psaB* gene to the *groEL* gene is inverted in the Lepto-03 and Lepto-04 lineage.

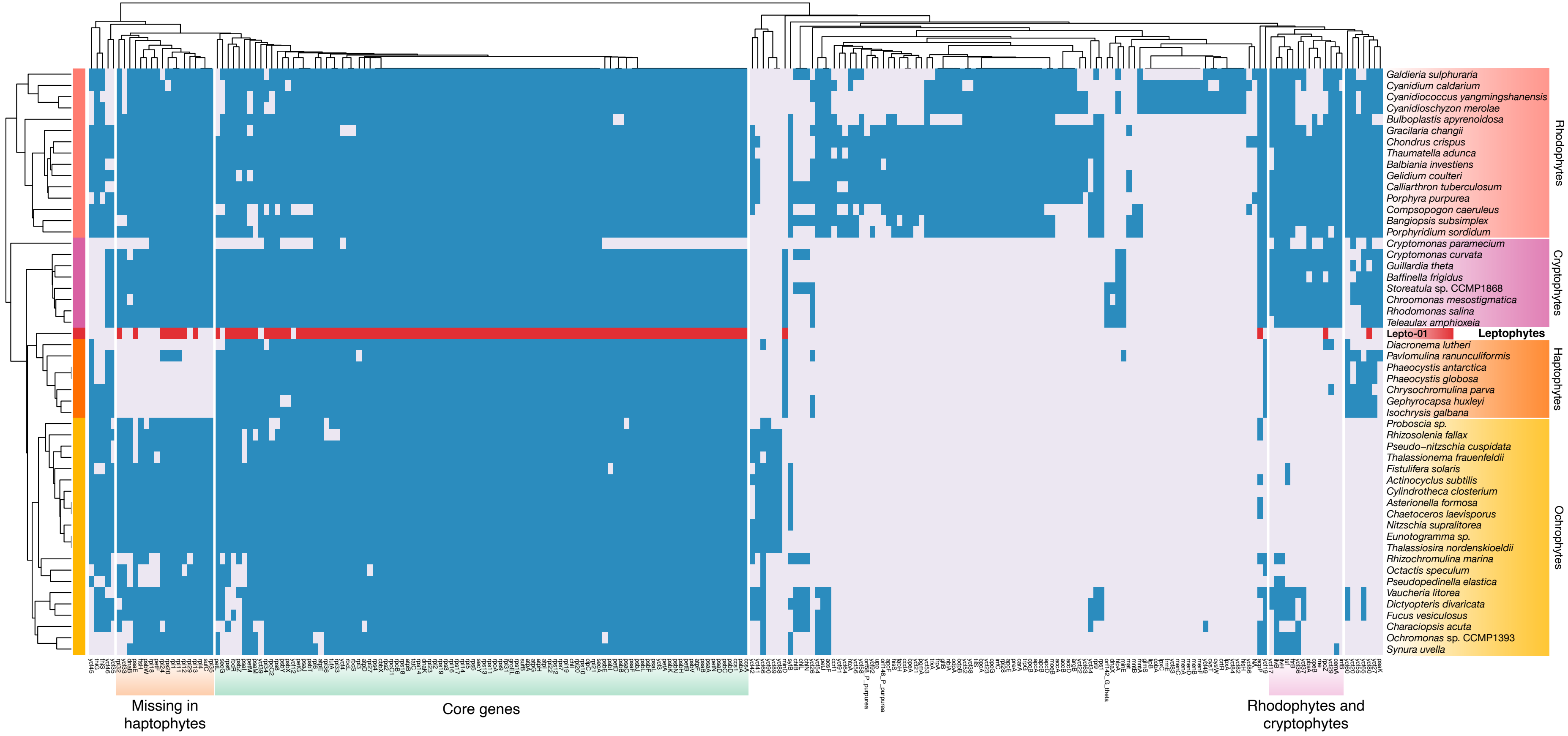


**Supplementary Fig. 17.** Binary heatmap and dendrograms of genes and taxa generated with UPGMA clustering, based on gene presence/absence of 237 plastid protein-coding genes. Blue/red and grey boxes represent presence and absence respectively.

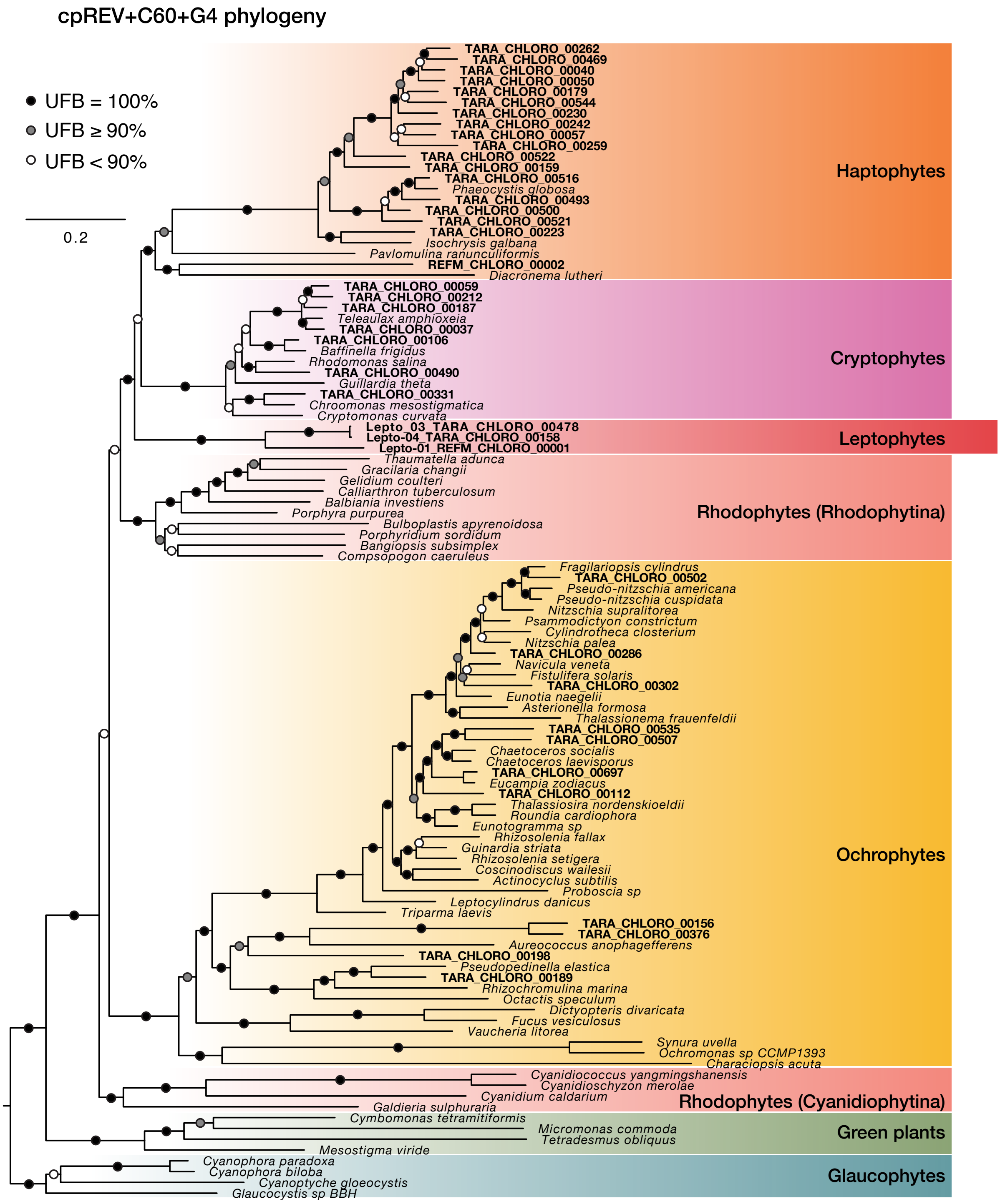


**Supplementary Fig. 18.** Maximum Likelihood phylogeny inferred with an alignment of 93 plastid-encoded genes (107 taxa, 20,292 sites). The phylogeny was inferred with IQ-TREE under the cpREV+C60+G4 model along with 1,000 ultrafast bootstraps.

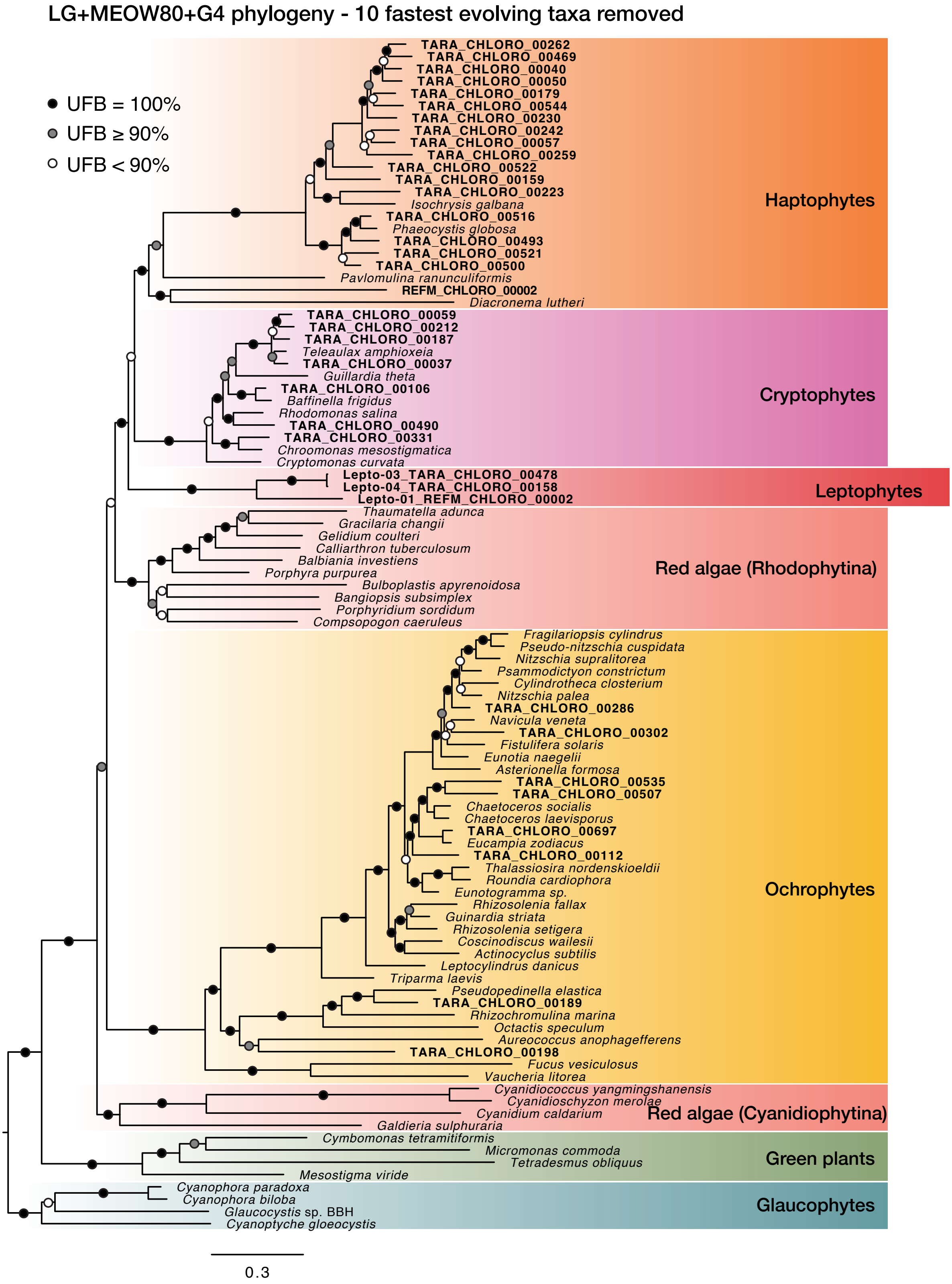


**Supplementary Fig. 19.** Maximum Likelihood phylogeny inferred with an alignment excluding the 10 fastest evolving taxa (longest root to tip distances). The input alignments included 93 plastid-encoded genes, 97 taxa, and 20,292 sites. The phylogeny was inferred with IQ-TREE under the LG+MEOW80+G4 model along with 1,000 ultrafast bootstraps.

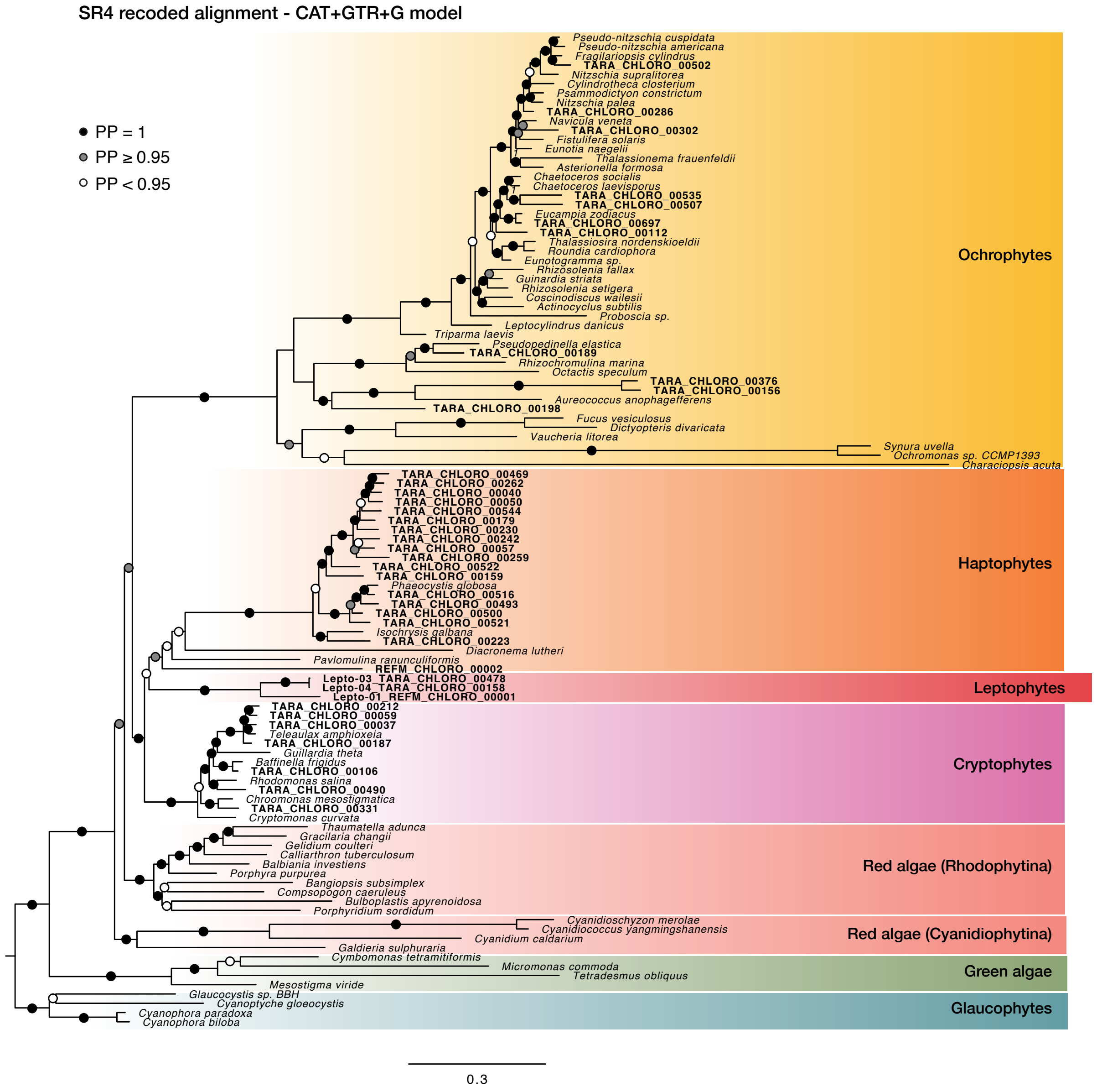


**Supplementary Fig. 20.** Bayesian inference of the SR4 recoded alignment with the CAT+GTR+G model (107 taxa, 20,292 sites).

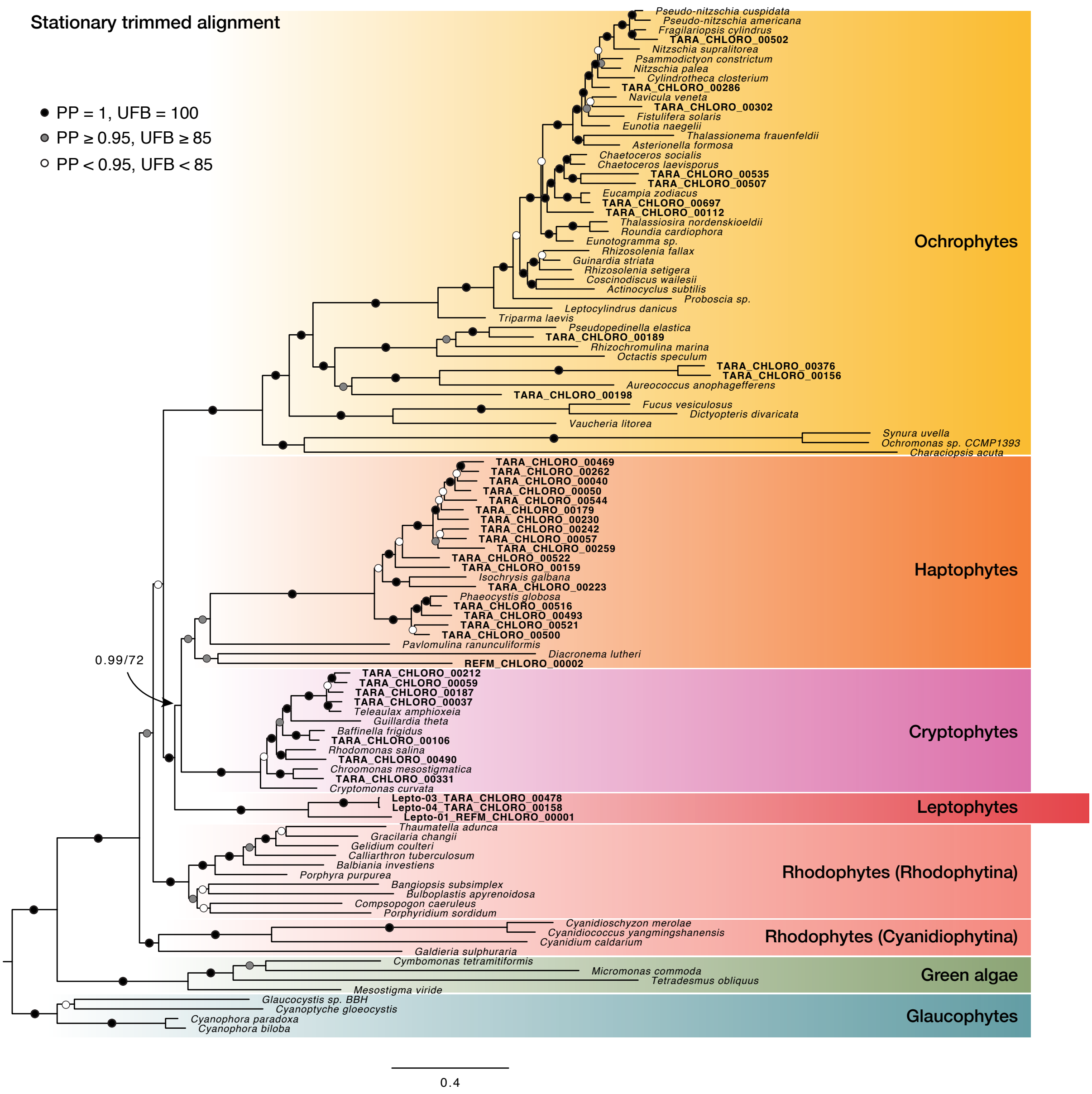


**Supplementary Fig. 21.** Bayesian inference of the stationary trimmed alignment to remove compositionally heterogenous sites (107 taxa, 15,754 sites). Branch support values are listed in the following order: posterior probabilities under the CAT+GTR+G model, and ultrafast bootstrap support under the LG+MEOW80+G model.

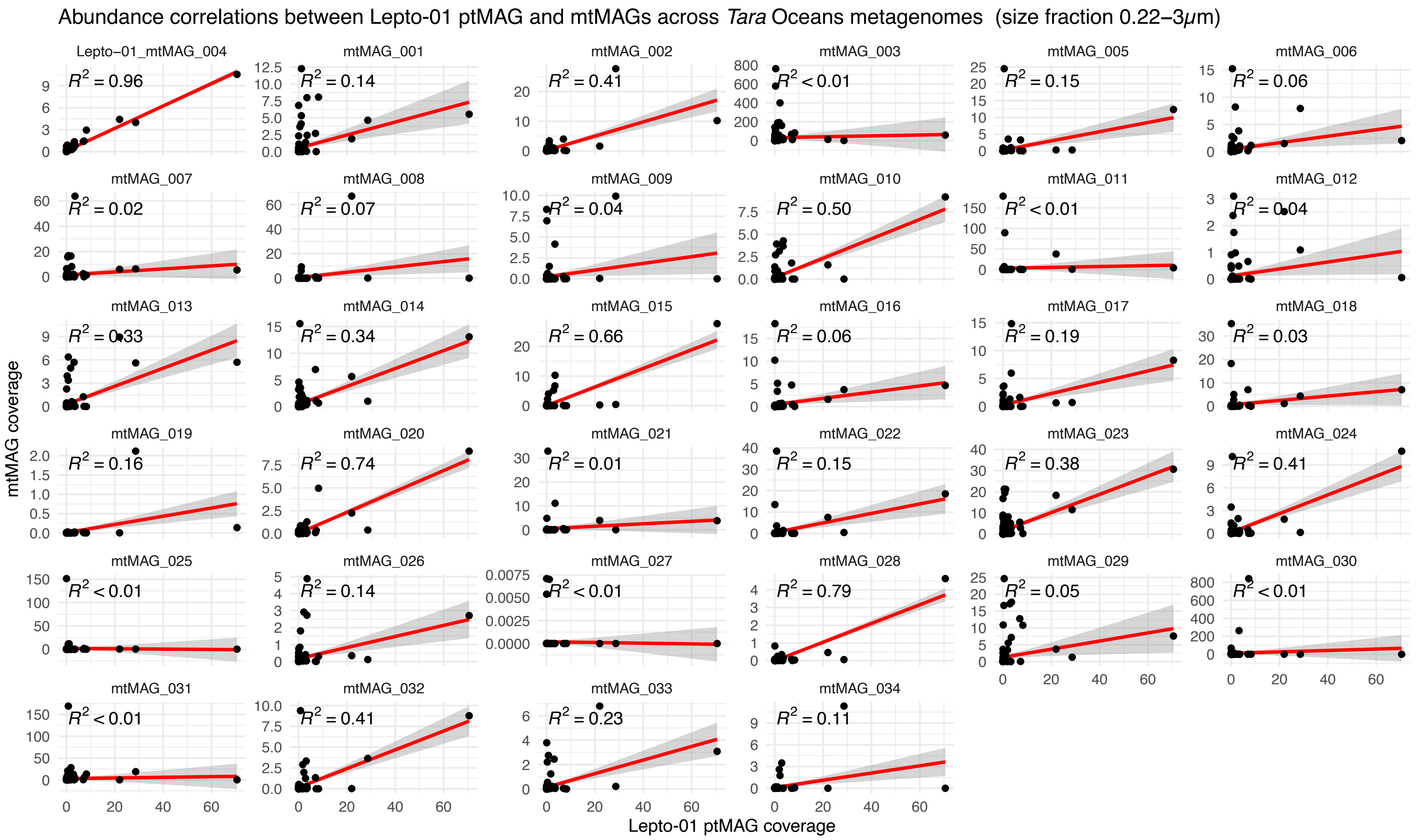


**Supplementary Fig. 22.** Abundance correlations between the Lepto-1 ptMAG and all 34 mtMAGs in Figure 4 across *Tara* Ocean metagenomes.

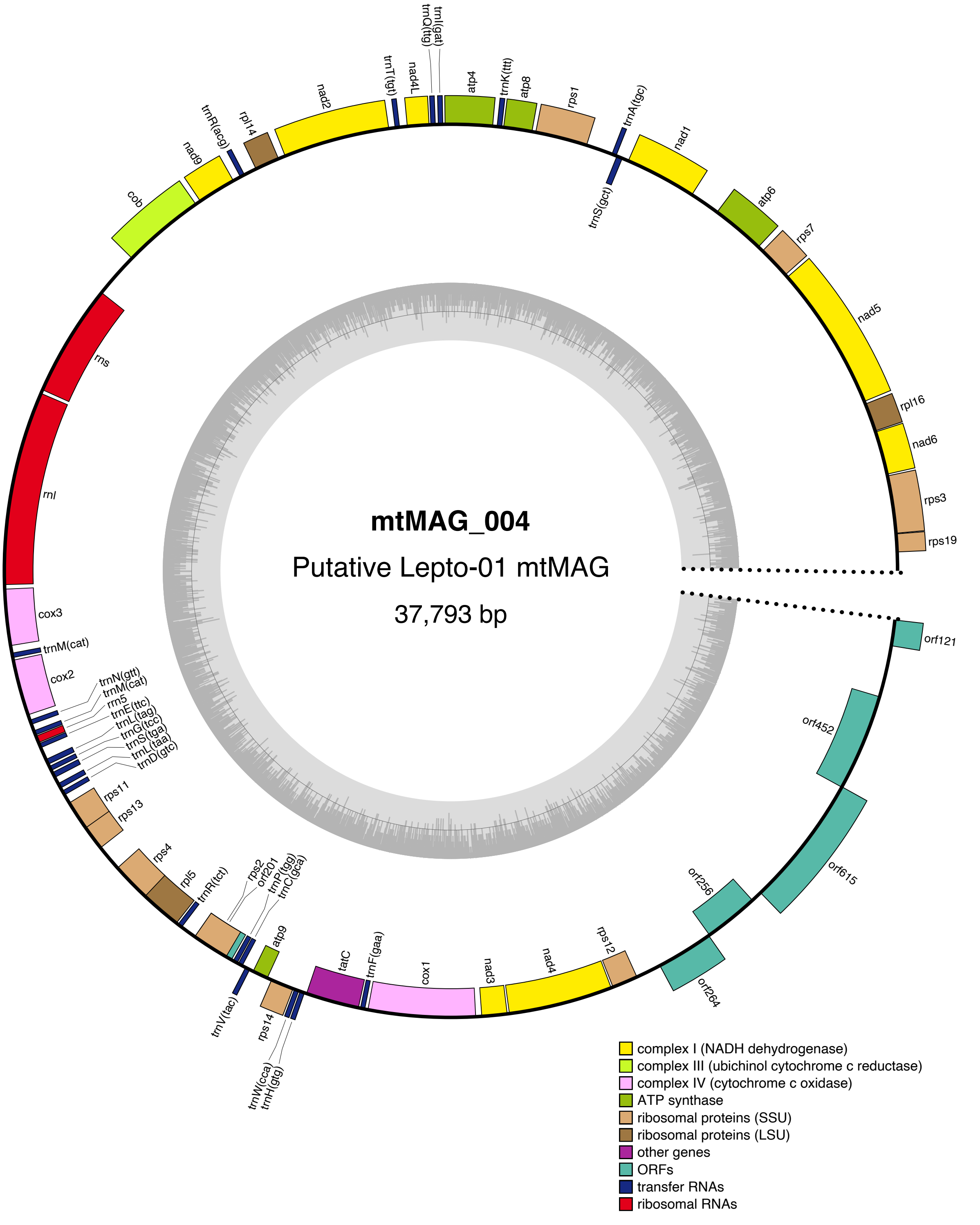


**Supplementary Fig. 23.** Annotated map of the putative Lepto-01 mitochondrial genome. Genes drawn inside the outer circle are transcribed clockwise, while those drawn outside the outer circle are transcribed counterclockwise. Genes are colour coded based on functional groups. The dark shaded area in the inner circle indicates GC content. The genome map was generated by OGDRAW[1].

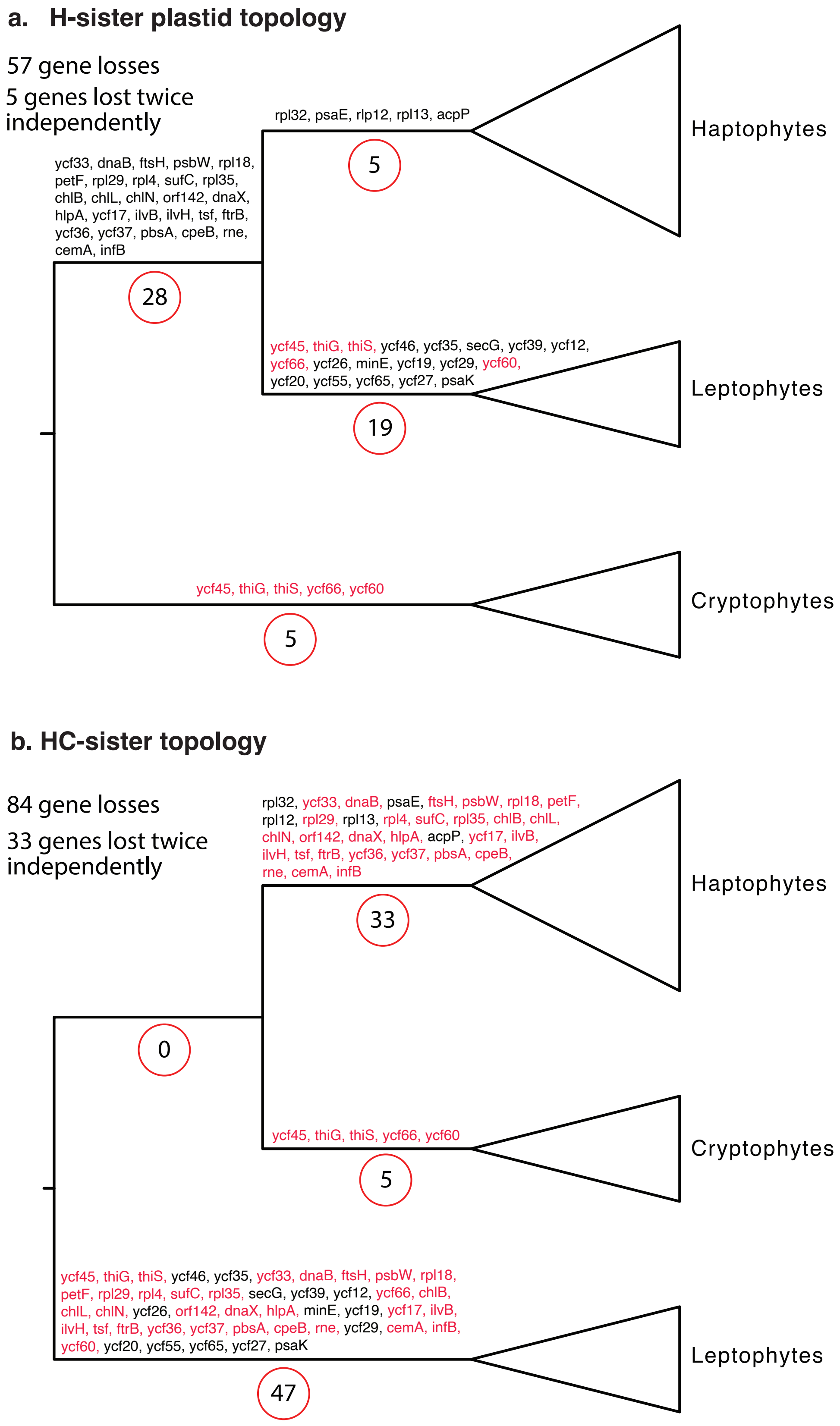


**Supplementary Fig. 24.** Maximum parsimony scenarios for gene losses under the two possible plastid topologies: (**a**) H-sister and (**b**) HC-sister. Genes lost multiple times independently are coloured red. Gene loss numbers (displayed in red circles at every branch) were calculated based on the taxon sampling in Fig. 2 and Supplementary Fig. 16.

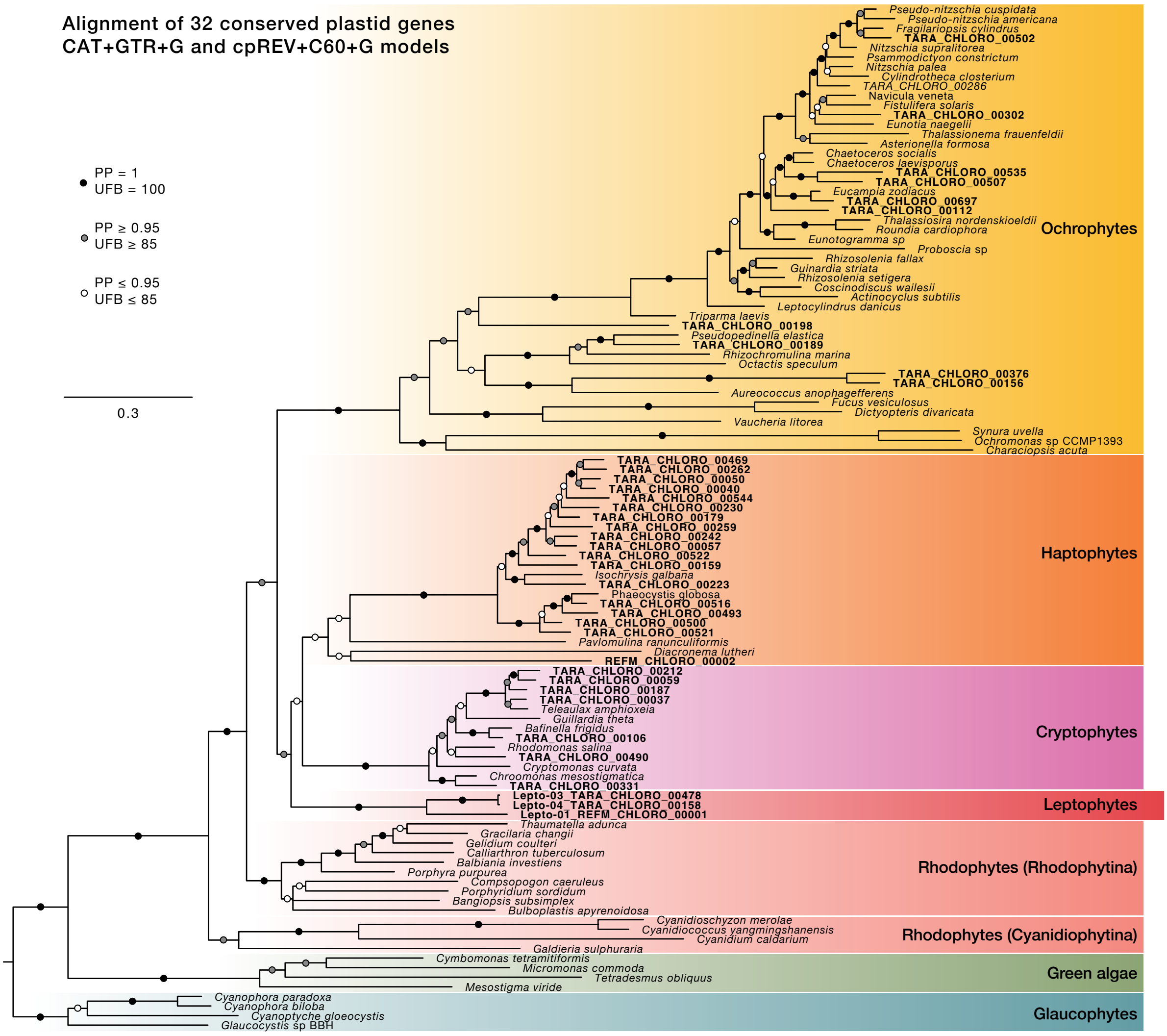


**Supplementary Fig. 25.** Bayesian inference of a preliminary dataset with 107 taxa, and 32 genes (7,921 sites). Branch support values indicate posterior probability obtained with the CAT+GTR+G model and ultrafast bootstrap values obtained with the cpREV+C60+G model.

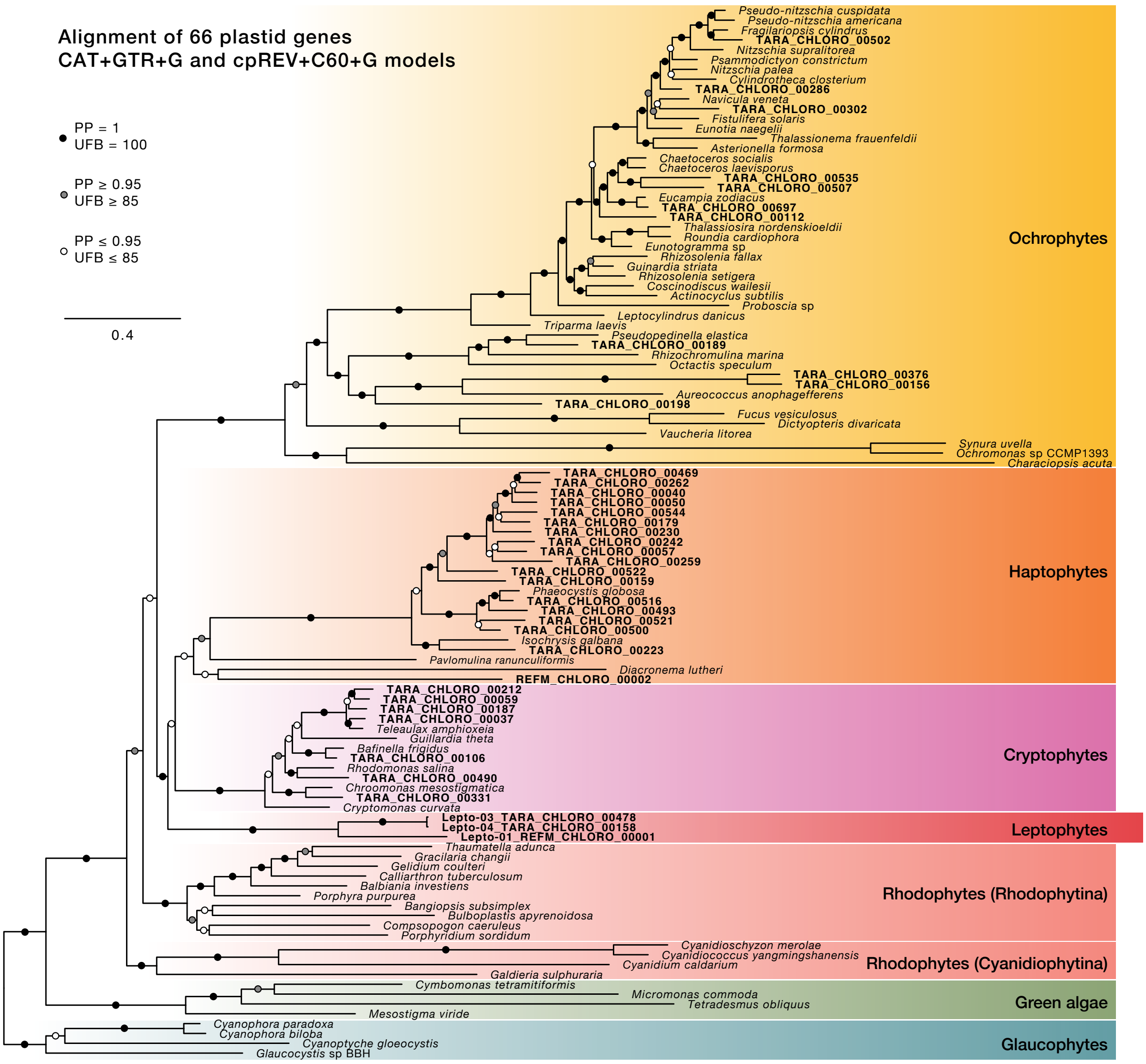


**Supplementary Fig. 26.** Bayesian inference of a preliminary dataset with 107 taxa, and 66 genes (16,438 sites). Branch support values indicate posterior probability obtained with the CAT+GTR+G model and ultrafast bootstrap values obtained with the cpREV+C60+G model.

**Supplementary Table 1.** (Provided as a separate excel file). Statistics of non-redundant ptMAGs and references and mapping results against *Tara* Ocean metagenomes.

**Supplementary Table 2.** Top 15 most abundant plastid genomes in the two Arctic stations where Lepto-01 (in bold text) is most abundant (0.8-2000 $\mu$m size fraction). Mean coverage was calculated by mapping the two listed metagenomes to each genome, and percentage abundance reflects each genome's signal as a proportion of the total plastid signal in each sample.

| | | | | | | | | |
|---|---|---|---|---|---|---|---|---|
| Region | **Arctic** | | | | **Arctic** | | | |
| Size fraction | **0.8-2000 µm** | | | | **0.8-2000 µm** | | | |
| Depth | **SUR** | | | | **SUR** | | | |
| Station | **194** | | | | **193** | | | |
| Filter ID | **194SUR1GGZZ11** | | | | **193SUR1GGZZ11** | | | |
| | **Plastid genome** | **Clade** | **Mean coverage** | **% abundance** | **Plastid genome** | **Clade** | **Mean coverage** | **% abundance** |
| | TARA_CHLORO_00329 | Ochrophyta (Coscinodiscophyceae) | 417.3643034 | 16.05 | NC_027589 | Cryptophyta (Teleaulax amphioxeia) | 514.6636486 | 25.84 |
| | TARA_CHLORO_00704 | Chlorophyta (Mammielophyceae) | 254.054665 | 9.77 | TARA_CHLORO_00704 | Chlorophyta (Mammielophyceae) | 306.51312 | 15.39 |
| | TARA_CHLORO_00272 | Euglenozoa | 186.9111833 | 7.19 | TARA_CHLORO_00237 | Chlorophyta (Mammielophyceae) | 153.864608 | 7.72 |
| | TARA_CHLORO_00680 | Ochrophyta (Coscinodiscophyceae) | 154.8617982 | 5.95 | NC_012575 | Chlorophyta (Micromonas commoda) | 83.05775298 | 4.17 |
| | TARA_CHLORO_00275 | Cryptophyta (Cryptophyceae) | 153.357879 | 5.90 | TARA_CHLORO_00297 | Ochrophyta (Coscinodiscophyceae) | 65.48039179 | 3.29 |
| | **Lepto-01** | **Leptophyte** | 124.151819 | 4.77 | TARA_CHLORO_00473 | Haptophyta (Prymnesiophyceae) | 62.23567659 | 3.12 |
| | TARA_CHLORO_00327 | Ochrophyta (Coscinodiscophyceae) | 88.50332919 | 3.40 | TARA_CHLORO_00686 | Ochrophyta | 53.42210228 | 2.68 |
| | TARA_CHLORO_00297 | Ochrophyta (Coscinodiscophyceae) | 84.94463191 | 3.27 | TARA_CHLORO_00326 | Ochrophyta (Coscinodiscophyceae) | 44.74423352 | 2.25 |
| | TARA_CHLORO_00473 | Haptophyta (Prymnesiophyceae) | 82.07171127 | 3.16 | TARA_CHLORO_00327 | Ochrophyta (Coscinodiscophyceae) | 43.57903498 | 2.19 |
| | TARA_CHLORO_00240 | Haptophyta (Prymnesiophyceae) | 75.97484423 | 2.92 | TARA_CHLORO_00240 | Haptophyta (Prymnesiophyceae) | 42.20396394 | 2.12 |
| | TARA_CHLORO_00328 | Ochrophyta (Coscinodiscophyceae) | 74.68082057 | 2.87 | TARA_CHLORO_00299 | Ochrophyta (Bacillariophyceae) | 33.79422825 | 1.70 |
| | TARA_CHLORO_00326 | Ochrophyta (Coscinodiscophyceae) | 70.60133521 | 2.71 | TARA_CHLORO_00282 | Ochrophyta (Bacillariophyceae) | 33.1461306 | 1.66 |
| | TARA_CHLORO_00263 | Haptophyta (Prymnesiophyceae) | 52.65510577 | 2.02 | **Lepto-01** | **Leptophyte** | 32.3916202 | 1.63 |
| | NC_012575 | Chlorophyta (Micromonas commoda) | 41.60626851 | 1.60 | TARA_CHLORO_00328 | Ochrophyta (Coscinodiscophyceae) | 29.42184233 | 1.48 |
| | TARA_CHLORO_00239 | Haptophyta (Prymnesiophyceae) | 41.37302726 | 1.59 | TARA_CHLORO_00267 | Ochrophyta (Dictyochophyceae) | 26.61431827 | 1.34 |

**Supplementary Table 3.** (On following page). Companion table for Figure 3c in the main text comparing six alternate topologies. Panel **a** shows the different topologies tested, focusing on three possible positions for leptophytes (HC-sister, H-sister, C-sister) under scenarios where complex plastids are monophyletic (as recovered in [4]) or non-monophyletic (as recovered in [5]). The table extends Figure 3c which displayed results only for the topologies where complex plastids are monophyletic for the sake of clarity. Panel **b** presents maximum likelihood estimates under different models for alternate positions of leptophytes within the CHL clade. The highest scoring topology is highlighted in bold. Blue bars show the difference in log-likelihood scores between each topology and the best-scoring one. A single asterisk (*) indicates topologies rejected by the Bonferroni-corrected chi-squared test, while two asterisks (**) indicate topologies also rejected by an Approximately Unbiased (AU) test. P-values are shown in Supplementary Table 4.

**a**

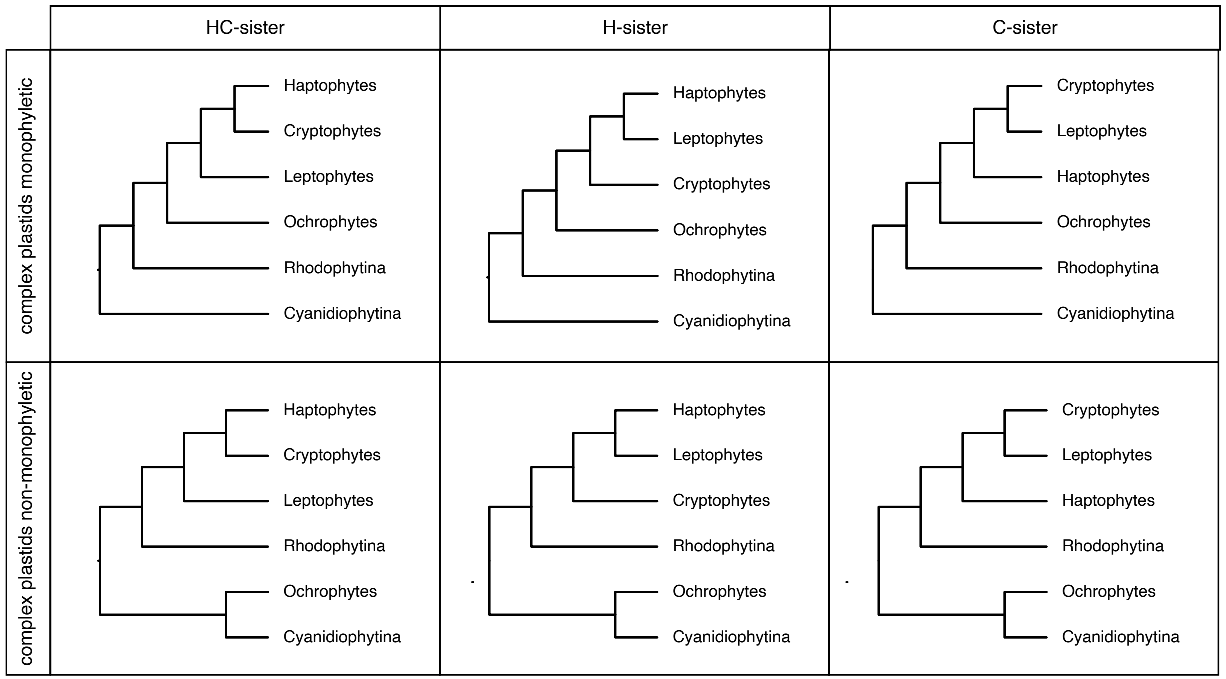


**b**

ML analyses using fixed topology trees

| **Dataset** | Untreated | | | | Stationary trimmed |
|---|---|---|---|---|---|
| **Model** | GTR-CAT-PMSF | LG-MEOW80 | LG-MEOW80 + GHOST (8 cat) | LG-MEOW80 + GF-MIX † | LG-MEOW80 |
| Topology | | | | | |
| HC-sister, complex plastids monophyletic | **-931909.236** | **-1056271.087** | **-1045458.696** | -1055620.569 * | **-592787.09** |
| H-sister, complex plastids monophyletic | -931920.074 * | -1056274.766 | -1045464.078 * | **-1055586.193** | -592793.164 * |
| C-sister, complex plastids monophyletic | -931924.826 ** | -1056273.024 | -1045463.152 * | -1055602.069 * | -592795.016 * |
| HC-sister, complex plastids non-monophyletic | -931915.342 | -1056272.398 | -1045466.547 * | -1055607.982 * | -592794.338 * |
| H-sister, complex plastids non-monophyletic | -931929.606 ** | -1056281.817 * | -1045478.6 * | -1055617.997 * | -592801.836 * |
| C-sister, complex plastids non-monophyletic | -931935.101 ** | -1056291.179 * | -1045482.379 * | -1055651.186 * | -592806.39 * |

* Rejected by Bonferroni corrected χ2 test

** Rejected by Bonferroni corrected χ2 test and AU test

† AU test not possible under GF-MIX model

**Supplementary Table 4.** (Provided as separate excel file). Support for alternative placement of leptophytes under different models with corresponding p-values for the Bonferroni corrected chi-squared test and the approximately unbiased (AU) test.

**Supplementary Table 5.** (Provided as separate excel file). Statistics of the 34 mtMAGs recovered from the co-assembly of six metagenomes where Lepto-01_REFM_CHLORO_00001 was most abundant.

**Supplementary Table 6.** (Provided as separate excel file). The 44 plastid encoded genes used to estimate completeness and redundancy (results show in Supplementary Figure 2 and Supplementary Table 1).

**Supplementary Table 7.** (Provided as separate excel file). The 25 canonical mitochondrial genes used to check for mitochondrial contamination in the plastid MAGs.

**Supplementary Table 8.** (Provided as separate excel file). The 93 plastid encoded genes used for phylogenomic analyses.

**Supplementary Table 9.** Convergence of Phylobayes GTR+CAT+G4 analyses for the 107 taxon, 93 gene dataset with a fixed input tree for the CAT-PMSF approach[6]. Two Markov chains were run, and burn-in for tracecomp was set to 1,800.

| Parameter | Effective sample size | Relative difference |
|---|---|---|
| loglik | 35 | 1.6552 |
| length | 113 | 0.0624932 |
| alpha | 330 | 0.525332 |
| Nmode | 146 | 0.227368 |
| statent | 118 | 3.53681 |
| statalpha | 110 | 0.210322 |
| rrent | 104 | 0.979222 |
| rrmean | 3525 | 0.0110885 |

**Supplementary Table 10.** (provided as an excel file). Hierarchical orthogroups (HOGs) identified by OrthoFinder used to generate Fig. 2 in the main text. HOGs were manually checked by inspecting gene annotations, single-gene trees, and performing BLAST and InterProScan searches.

**Supplementary Table 11.** (provided as an excel file). Taxa (along with accession numbers) used for mitochondrial phylogenies (Figure 4 of main text).

**Supplementary Table 12.** (provided as an excel file). Genes used for mitochondrial phylogenies (Figure 4 of main text).

### Supplementary Note 1.
### Evaluating the position of leptophytes in the plastid phylogeny

Our phylogenomic analyses based on 107 taxa, 93 plastid-encoded genes and a range of sophisticated phylogenetic models confirmed that leptophytes are related to haptophytes and cryptophytes. However, the position of leptophytes within the CHL group (Cryptophytes-Haptophytes-Leptophytes) could not be confidently resolved and two topologies remained credible: the HC-sister toplogy (leptophytes sister to both hatophytes and cryptophytes), and the H-sister topology (leptophytes sister to haptophytes).

The uncertainty of the position of leptophytes could be caused by lack of phylogenetic signal (as indicated by the small branch lengths separating the groups within the CHL clade), and could also be caused by potential biases and phylogenetic artefacts, such as potential compositional heterogeneity across taxa[1], and heterotachy[2].

The LG+MEOW80+GHOST model with 8 linked site classes supported the HC-sister topology, which was also recovered by the CAT-GTR+G model with strong support (Fig. 3 in main text). However, the GFmix[1] model which accounts for compositional heterogeneity across sites and branches strongly favoured the H-sister topology (maximum likelihood values for the H-sister and HC-sister topology differed by 34 points).

##### Compositional heterogeneity

Compositional heterogeneity across taxa may be an artefact of our dataset, particularly due to the inclusion of extremophile red algae (Cyanidiophytina). Additionally, recent studies have suggested that ribosomal protein genes, which are highly represented in our dataset, tend to exhibit greater compositional bias due to strong co-evolution among these proteins[3,4]. A principal component analysis (PCA) revealed the amino acid preferences across different groups (Supplementary Fig. 27).

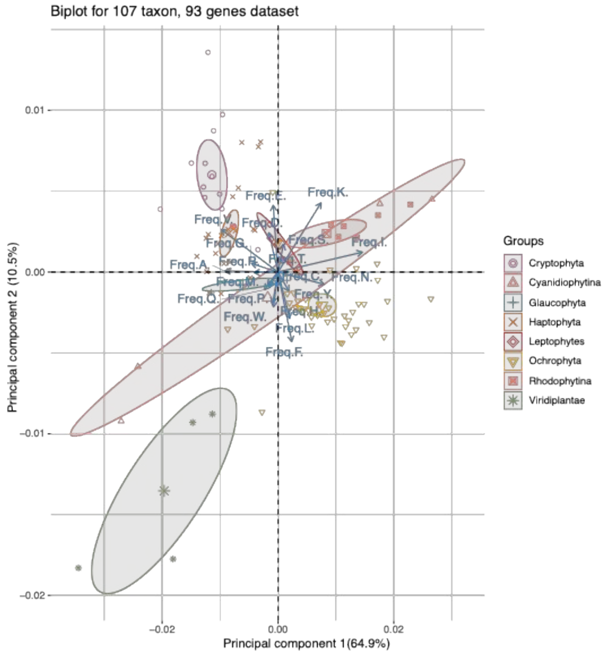


**Supplementary Figure 27.** PCA analysis showing compositional biases among clades based on the alignment used for phylogenomic analyses.

We accounted for the compositional bias in three ways: (1) evaluating the likelihood of alternative topologies under the GFmix model, (2) removing compositionally heterogenous sites, and (3) recoding the dataset using the SR4 recoding scheme[5].

1. *GFmix model*

The GFmix model adjusts the vector of amino acid frequencies in a branch-specific manner to account for shifts in the relative frequencies of amino acids in different branches of the tree. To identify which amino acids are enriched or depleted in specific lineages, we analyzed our concatenated alignment using a custom R script (courtesy of Charley McCarthey, Dalhousie University; available on the GitHub repository: https://github.com/burki-lab/beta-Cyclocitral). This script divided the taxa into two groups and identified amino acid groups enriched or depleted in each lineage. The two taxa groups compared were *Cyanidiococcus yangmingshanensis* (accession NC_051883) and *Cyanidioschyzon merolae* (accession NC_004799) versus all other taxa.

The groups of amino acids enriched or depleted in these lineages are as follows:
**G-class**: I, N, F, K, T, D, S, E
**F-class**: R, C, V, W, H, M, A, Q

The GFmix program utilized the enriched and depleted amino acid classes to calculate the log-likelihoods for the six alternative topologies listed in Supplementary Table 4a. Among these, the H-sister topology with complex plastids forming a monophyletic grouping, was strongly favoured by the LG+MEOW80+GFmix model (Supplementary Table 4a). All other topologies were statistically rejected based on the Bonferroni-corrected chi-squared test.

2. *Removing compositionally heterogenous sites*

We generated a compositionally homogenized dataset by removing heterogenous sites from the concatenated alignment using Stuart's test of marginal homogeneity as implemented in BMGE[6]. This dataset was then analysed with the conventional site-heterogenous model LG+MEOW80+G, recovering the HC-sister topology (Fig. 3c, Supplementary Table 4). All alternative topologies were rejected by the Bonferroni-corrected chi-squared test, but not by the AU topology test. The HC-sister topology was also strongly supported when analysing the dataset under the CAT+GTR+G model (PP = 0.99).

3. *SR4 recoding*

We recoded the concatenated alignment using the SR4 recoding scheme and inferred a phylogeny under the CAT+GTR+G model in PhyloBayes. This phylogeny also supported the H-sister topology, consistent with the results of the GFmix model, albeit with low support (PP = 0.75). The limited support may stem from the low phylogenetic signal in the dataset, further eroded by amino acid recoding. Additionally, we caution that this result should be interpreted cautiously as recoding datasets has been shown to generate incorrect trees due to the associated of phylogenetic signal[7,8].

Altogether, the GFmix model, which explicitly models both site- and branch-heterogeneity, supports the H-sister topology. However, compositionally homogenized datasets analysed with conventional site-heterogenous models recapitulate the results of the GFmix model in only one out of two cases. While H-sister topology is more likely as it invokes a simpler scenario of plastid transfer and gene content evolution (Fig. 5, Supplementary Fig. 24), we cannot confidently rule out the HC-sister topology based on our current analyses. More data in the form of cell morphology and nuclear DNA from leptophytes will be required to decide between these scenarios.

**References:**


1. Muñoz-Gómez, S. A. *et al.* Site-and-branch-heterogeneous analyses of an expanded dataset favour mitochondria as sister to known Alphaproteobacteria. *Nat. Ecol. Evol.* **6**, 253–262 (2022).
2. Crotty, S. M. *et al.* GHOST: Recovering Historical Signal from Heterotachously Evolved Sequence Alignments. *Syst. Biol.* **69**, 249–264 (2020).
3. Petitjean, C., Deschamps, P., López-García, P. & Moreira, D. Rooting the Domain Archaea by Phylogenomic Analysis Supports the Foundation of the New Kingdom Proteoarchaeota. *Genome Biol. Evol.* **7**, 191–204 (2014).
4. Ramulu, H. G. *et al.* Ribosomal proteins: Toward a next generation standard for prokaryotic systematics? *Mol. Phylogenet. Evol.* **75**, 103–117 (2014).
5. Susko, E. & Roger, A. J. On Reduced Amino Acid Alphabets for Phylogenetic Inference. *Mol. Biol. Evol.* **24**, 2139–2150 (2007).
6. Criscuolo, A. & Gribaldo, S. BMGE (Block Mapping and Gathering with Entropy): a new software for selection of phylogenetic informative regions from multiple sequence alignments. *BMC Evol. Biol.* **10**, 210 (2010).
7. Hernandez, A. M. & Ryan, J. F. Six-State Amino Acid Recoding is not an Effective Strategy to Offset Compositional Heterogeneity and Saturation in Phylogenetic Analyses. *Syst. Biol.* **70**, 1200–1212 (2021).
8. Foster, P. G. *et al.* Recoding Amino Acids to a Reduced Alphabet may Increase or Decrease Phylogenetic Accuracy. *Syst. Biol.* **72**, 723–737 (2023).
